# Discovery of oncogenic ROS1 missense mutations with sensitivity to tyrosine kinase inhibitors

**DOI:** 10.1101/2022.12.27.521482

**Authors:** Sudarshan R. Iyer, Kevin Nusser, Kristen Jones, Pushkar Shinde, Catherine Z. Beach, Clare Keddy, Erin Aguero, Jeremy Force, Ujwal Shinde, Monika A. Davare

## Abstract

Chromosomal rearrangements of *ROS1* generate ROS1 tyrosine kinase fusion proteins that are established oncogenes predicting effectiveness of tyrosine kinase inhibitors (TKI) treatment. The cancer genome reveals nonsynonymous missense mutations in *ROS1,* however, their oncogenic potential remains unknown. We nominated thirty-four tumor-associated missense mutations in ROS1 kinase domain for functional interrogation. Immunoblotting revealed diverse impact of the mutations on the kinase, ranging from loss of function to significant increase in catalytic activity. Notably, Asn and Gly substitutions at the Asp-2113 position in ROS1 kinase domain were TKI- sensitive hyper-activating mutations, and transformative oncogenes in independent cell models. Molecular modeling revealed drastic alterations in the activation loop of ROS1^D2113N^ compared to wildtype kinase. Proteomics studies showed that ROS1^D2113N^ increases phosphorylation of known effectors akin to ROS1 fusions, and upregulates pathways not previously linked to ROS1, including mTORC2, JNK1/2, AP-1, TGFB1 and CCN1/2. *In vivo*, ROS1^D2113N^ drove tumor formation that was sensitive to inhibition by crizotinib and lorlatinib. Taken together, these data show that select point mutations within ROS1 RTK are oncogenic, and maybe therapeutically targetable with FDA-approved TKI.

## Introduction

Targeted therapy of cancers driven by aberrant receptor tyrosine kinase (RTK) signaling has achieved remarkable clinical outcomes over the past two decades (Drilon *et al*, 2021; Schram *et al*, 2017). RTKs normally serve to link extracellular signals to intracellular signaling pathways to influence various cell-based phenotypes such as, but not limited to, cell proliferation, differentiation, and changes in metabolism (Lemmon & Schlessinger, 2010). In the case of cancer, these proteins become aberrantly activated, which leads to upregulation of several downstream signaling pathways where the net result becomes uncontrolled cell proliferation and survival (Du & Lovly, 2018). To combat such molecular aberrations, tyrosine kinase inhibitors (TKIs) have been developed to abrogate kinase activity and thus shut off constitutive upregulation of pro-cell proliferation signaling programs. TKIs have shown robust efficacy in reducing cancer burden in patients as detailed in many clinical trials (Cocco *et al*, 2018; Drilon *et al*., 2021; Drilon *et al*, 2020; Landi *et al*, 2019).

Diverse germline or somatic aberrations result in a gene gaining oncogene characteristics that drive malignant transformation. Among these, gain-of-function, nonsynonymous point mutations such as EGFR^L858R^ have been established to confer constitutive catalytic activity and induce oncogenic signaling (Brewer *et al*, 2013; Sharma *et al*, 2007). With widespread adoption of targeted next-generation sequencing (NGS) of tumors in the clinical setting, as well as the generation of large datasets from previous research cohorts by The Cancer Genome Atlas Project, many novel somatic variants in known oncogenes have been uncovered. However, the majority of these are still variants of unknown significance (VUS), i.e., it is unknown if a certain variant contributes to tumor growth versus if it merely serves as a ‘passenger’ without functional impact on tumor biology. This represents a significant bottleneck for clinical translation using NGS data. Various *in silico* algorithms that attempt to predict the effect of mutations on protein structure and function exist (Reva *et al*, 2007, 2011; Sim *et al*, 2012; Thusberg & Vihinen, 2009; Vaser *et al*, 2016), yet in our experience, they are neither accurate nor reliable enough to replace functional validation in a lab setting. An approach that involves an initial *in silico* prediction step to filter variants to a smaller and more feasible list of variants for laboratory-based functional testing may facilitate efficient discovery of new oncogenic, gain-of-function mutations amenable for targeted therapy.

Like other RTKs, the proto-oncogene *ROS1*, has shown an ability to promote oncogenic transformation when aberrantly activated (Drilon *et al*., 2021). To date, the role of *ROS1* in cancer is largely restricted to the oncogenic ROS1 fusion proteins resulting from chromosomal rearrangements. These ROS1 fusion oncogenes harbor unchecked constitutive catalytic activities resulting from the loss of the regulatory amino-terminal domain of the receptor. ROS1 fusion proteins are established oncogenic drivers in a diverse set of cancers, with the most extensive body of literature coming from lung adenocarcinomas (Arai *et al*, 2013; Inoue *et al*, 2016; Lin & Shaw, 2017; Saborowski *et al*, 2013). Previous preclinical studies established that oncogenic ROS1 fusion proteins are effectively targeted with tyrosine kinase inhibitors (TKIs) (Davare *et al*, 2013; Davies *et al*, 2012). Subsequently, robust clinical responses were achieved with the TKIs crizotinib and entrectinib in ROS1-fusion-positive non-small cell lung cancer (NSCLC), which led to FDA approval (Drilon *et al*, 2017b; Shaw *et al*, 2014). While ROS1 fusion proteins now have an established role as targetable oncogenes in cancer, whether *ROS1* missense mutations or amplifications can drive or contribute to tumor formation is unknown (Drilon *et al*., 2021). Such knowledge could potentially expand the cohort of cancer patients that may benefit from efficacious FDA-approved ROS1-TKI therapy.

In this study, we hypothesized that a subset of *ROS1*nonsynonymous missense mutations are gain-of-function variants that contribute to tumor formation and can be targeted with TKI treatment. To this end, we used the AACR Genie dataset for discovery of *ROS1* missense variants in a tumor-agnostic fashion and, based on *in silico* filtering strategies, nominated mutations that specifically reside in the tyrosine kinase domain for wet-lab characterization (Consortium, 2017; Sim *et al*., 2012; Vaser *et al*., 2016). Using biochemical approaches, we identified gain of function ROS1 mutations and performed *in vitro* and *in vivo* transformation experiments as well as structural modeling, molecular dynamics simulation, and global proteomics to validate novel oncogenic *ROS1* variants.

## Materials and Methods

### Cell lines, compounds, and plasmids

HEK-293T/17 (ATCC Cat# CRL-11268, RRID:CVCL_1926), MCF10A cells (ATCC Cat# CRL-10317, RRID:CVCL_0598), and NIH-3T3 cells (ATCC Cat# CRL-1658, RRID:CVCL_0594) were purchased from American Type Culture Collection (ATCC; Manassas, VA, USA). HEK-293A cells (Cat# R70507; RRID:CVCL_6910) was purchased from Thermo Fisher Scientific (Waltham, MA, USA). Platinum-A and Platinum-E cells were purchased from Cell BioLabs, Inc. (San Diego, CA, USA). Primary antibodies used in the study are as follows: phospho-ROS1 (Cell Signaling Technology, Inc. (CST; Danvers, MA, USA); Y2274; #3078; RRID:AB_2180473), ROS1 (CST; #3287; RRID:AB_2797603), phospho-STAT3 (CST; Y705; #9145; RRID:AB_2491009), STAT3 (CST; #9139; RRID:AB_331757), phospho-ERK (CST; T202/Y204; #9101; RRID:AB_331646), ERK (CST; #4696; RRID:AB_390780), anti-FLAG (Thermo Fisher Scientific Cat# 701629, RRID:AB_2532497), α-TUBULIN (Developmental Studies Hybridoma Bank (DSHB; Iowa City, IA, USA); 12G10; RRID:AB_1157911), ACTIN (DSHB; JLA20; RRID:AB_528068), GAPDH (Santa Cruz Biotechnology, Inc.; Dallas, TX, USA; sc-47724; RRID:AB_627678), phospho-S6 (CST; S235/236, #4858; RRID:AB_916156), and S6 (CST; #2317; RRID:AB_2238583). DMEM, DME-F/12, L-glutamine, and antibiotics were purchased from Genesee Scientific (San Diego, CA, USA). Fetal bovine serum (FBS) was procured from Neuromics (CA3 Biosciences, Inc., Edina, MN, USA). Bovine growth serum (BGS) was procured from VWR International (Radnor, PA, USA). Horse serum (HS) and calf serum (CS) were purchased from Thermo Fisher Scientific. Bovine pancreas insulin solution was purchased from Millipore Sigma (Cat# I0516; Burlington, MA, USA). Recombinant human EGF was purchased from Peprotech (Cranbury, MA, USA), hydrocortisone powder from Millipore Sigma (Cat# H0888), and cholera toxin from Cayman Chemical (Ann Arbor, MI, USA). 0.05% Trypsin/0.53 mM EDTA was procured from Corning Inc. (Corning, NY, USA) while TrypLE™ was purchased from Thermo Fisher Scientific. Crizotinib, cabozantinib, and staurosporine were purchased from LC Laboratories® (Woburn, MA, USA), and lorlatinib, entrectinib, repotrectinib, and cycloheximide from Selleck Chemicals Inc. (Houston, TX, USA). TransIT-LT1 transfection reagent was purchased from Mirus Bio LLC (Madison, WI, USA). Gateway LR clonase was purchased from ThermoFisher. Protease and phosphatase inhibitors were purchased from Bimake (Houston, TX, USA). Pre-cast Bolt™ and NuPAGE™ 4-12% Bis-Tris protein gels and 4X LDS sample buffer were purchased from ThermoFisher. Pierce™ BCA Protein Assay Kit was purchased from ThermoFisher. CCK-8 reagent was purchased from Bimake. (Milpitas, CA, USA). Infusion Cloning HD Kit was procured from Takara Bio USA Inc. (San Jose, CA, USA). pCMV-VSV-G was a gift from Bob Weinberg (Addgene plasmid # 8454; http://n2t.net/addgene:8454; RRID:Addgene_8454) (Stewart *et al*, 2003), pENTR4-FLAG (w210-2) was a gift from Eric Campeau & Paul Kaufman (Addgene plasmid # 17423 ; http://n2t.net/addgene:17423 ; RRID:Addgene_17423), pGP (retroviral Pol and Rev gene plasmid) was a gift from Romel Somwar, and pCX4-puro was a gift from Tsuyoshi Akagi. Wild- type full-length ROS1 human cDNA was purchased from DNASU (Tempe, AZ, USA).

Oligonucleotides were ordered from Integrated DNA Technologies Corp. (Newark, NJ, USA) and Eurofins Genomics (Eurofins Scientific, Luxembourg).

### Growth and propagation of cell lines

All cell lines were grown in a tissue culture incubator with 5% CO2 at 37°C. Cells were maintained in 75 cm^2^ flasks and subcultured when approaching 75% confluence. NIH-3T3 cells were maintained in DMEM medium supplemented with 10% (vol/vol) CS and 1% (vol/vol) L- glutamine solution and detached using 0.05% Trypsin/0.53 mM EDTA. HEK-293T/17, HEK- 293A, Platinum-A, and Platinum-E cells were maintained in DMEM medium supplemented with 10% (vol/vol) BGS and 1% (vol/vol) L-glutamine solution and detached using 0.05% Trypsin/0.53 mM EDTA. MCF10A cells were maintained in DMEM/F12 medium supplemented with 5% (vol/vol) horse serum (HS), 1% (vol/vol) L-glutamine solution, 0.5 mg/mL hydrocortisone, 100 ng/mL cholera toxin, 10 µg/mL insulin, and 20 ng/mL EGF (MCF10A growth medium) as previously described (Isakoff *et al*, 2005). MCF10A cells were detached using 2X TrypLE™ in 1X Versene and neutralized with MCF10A resuspension medium (DMEM/F12 supplemented with 20% (vol/vol) HS and 1% (vol/vol) L-glutamine solution). All cell lines were routinely tested for mycoplasma contamination (Lonza MycoAlert^TM^ PLUS Mycoplasma Detection Kit) and verified to be free of contamination. Antibiotics were included in all cell culture media.

### Cloning

The retroviral construct pCX4 ROS1 was cloned as described previously (Davare *et al*., 2013). Briefly, full-length ROS1 cDNA was cloned using the SalI and XhoI sites in the multiple cloning site of pENTR4-No ccDB (696-1) vector (Addgene Plasmid #17424) via In-Fusion™ cloning using PCR amplification that included the adition of C-terminal Flag tag . Simultaneously, pCX4- puro was converted into a Gateway™ Destination vector (pCX4-DEST-Puro) by inserting the Gateway™ Reading Frame Cassette A (ThermoFisher) into the multiple cloning site using EcoRI and XhoI restriction sites. ROS1-FLAG cDNA was subcloned into pCX4-DEST-Puro via LR clonase reaction. ROS1 mutants were generated using site-directed mutagenesis (QuikChange™ Mutagenesis Protocol, Agilent Technologies Inc., Santa Clara, CA, USA) of the pENTR4-ROS1- FLAG construct and subsequently subcloned into pCX4-DEST-Puro.

### Generation of stable isogenic cell lines

Replication incompetent, infectious ecotropic and VSV-G pseudotyped amphotropic retroviral particles were generated using Platinum-E and HEK293T/17 cells or Platinum A cells, respectively. Platinum-E cells were transfected in 6-well plates with 2 µg pCX4 ROS1 transfer plasmid complexed with 8 µL TransIT-LT1 reagent while HEK293T/17 cells were transfected in 10 cm dishes with 6 µg transfer plasmid, 5 µg pGP, and 4 µg pCMV-VSV-G complexed with 45 µL TransIT-LT1 reagent. Ecotropic viral supernatant collected at 48- and 72-hours post transfection was filtered with 0.45 µm syringe filter, and used for transduction of cells pre-treated with 2 µg/mL polybrene. VSV-G pseudotyped viral supernatant was concentrated via ultracentrifugation and used for cell transduction. Transduced cells were selected with 1 µg/mL puromycin for 4 days.

### Immunoblotting

Lysates were prepared from cells using a standard cell lysis buffer as described before (Davare *et al*, 2015). Protein quantitation was performed with the Pierce™ BCA Protein Assay Kit. For immunoblotting, we loaded 15 µg of reducing LDS sample buffer-extracted cleared cell lysates on pre-cast 4-12% Bolt™/NuPAGE™ Bis-Tris gels. Spectra Multicolor Broad Range Protein Ladder (Thermo Fisher Scientific) was used to determine relative molecular weights of protein bands after imaging. Proteins were transferred to nitrocellulose membranes and probed with indicated antibodies as recommended by the manufacturer. Western blots were imaged using the ChemiDoc™ (Bio-Rad Laboratories, Hercules, CA, USA) for detection of horseradish peroxidase-conjugated secondary antibodies or the Odyssey® DLx Imaging System (a LICOR, Lincoln, NE, USA) for detection of near-infrared fluorescent-conjugated secondary antibodies.

Phospho-ROS1 detection required the SuperSignal™ West Femto Maximum Sensitivity Substrate (Thermo Fisher Scientific). Densitometry was performed using Image Lab™ Software (RRID: SCR_014210).

### Transient transfection assays to screen ROS1 TKD mutants

HEK-293T/17 cells were grown to 70% confluence in 12-well standard tissue culture dishes, and transfected with 2 µg pCX4 ROS1 wildtype or mutant variants using 8 µL TransIT-LT1 reagent. After 48 h, lysates prepared from transfected cells were harvested and an equal volume of 10 µL was loaded on 4-12% NuPAGE™ Bis-Tris protein gels. Methods for immunobotting are described above. Relative ROS1 catalytic activity was determined using a ratio of densitomety data from phospho-ROS1 (pROS1) and total ROS1 (tROS1) antibody signals. Mutants showing higher pROS1/tROS1 ratio relative to wild-type were considered activating.

### Cell proliferation and oncogenic transformation assays

#### MCF10A proliferation assay

Stable MCF10A cells expressing ROS1 variants were seeded in 96 well plates and treated with DMSO or different ROS1-TKIs (n=5 per condition). For this assay, a modified growth medium was used where concentrations were reduced from 2ng/ml to 0.5ng/ml for EGF and 10% to 2% for HS, respectively. The final DMSO % was kept at ≤ 0.1%. 5000 cells were seeded in a final working volume of 200 µL. Cells were grown in these conditions for seven days. Real-time proliferation was monitored using the Incucyte® ZOOM imaging platform and final cell confluence was measured via the CCK-8 (WST-8, tetrazolium)-based cell viability assay per manufacturer’s protocol; 490 nm absorbance was read at 4 hours post addition of CCK- 8 reagent using a BioTek Synergy™ H1 plate reader and CCK-8 absorbance values of inhibitor- treated wells were normalized to vehicle-treated wells. Data analysis and graph generation was using GraphPad Prism v9.3 (GraphPad Software, RRID:SCR_002798, San Diego, CA, USA).

#### NIH-3T3 anchorage-independent soft agar growth assay

Anchorage-independent soft agar growth experiments were performed as described (Davare *et al*., 2013). Briefly, 8000 cells (stable NIH-3T3 ROS1 variant cell lines) were seeded in a 0.2% top agarose layer layered on top of a 0.4% bottom agar layer. As indicated inhibitors were added in the feeding medium after top matrix solidified for 24 h. Every week, for upto 4 weeks, colonies were counted using the Gelcount™ colony counter (Oxford Optronix Ltd., Milton Park, Abingdon, UK). Resulting data were analyzed with Microsoft Excel (Microsoft Excel, RRID:SCR_016137) and GraphPad Prism.

### Analysis of AACR Genie clinical sequencing data to find somatic ROS1 TKD mutations

AACR Genie data was queried for the *ROS1* gene and then downloaded from genie.cbioportal.org (Cerami *et al*, 2012; Gao *et al*, 2013). All datasets on the website, including data from GENIE Cohort v11.0-public, DFCI-Profile Glioma Cohort 2013-2018, AACR Project GENIE AKT1 Cohort, and Metastatic Breast Cancer 2013-2016 datasets were included (Consortium, 2017; Garrido-Castro *et al*, 2021; Touat *et al*, 2020). The full list of ROS1 aberrations was narrowed down further using the ‘Sorting Intolerant From Tolerant’ (SIFT) scores (Sim *et al*., 2012) and limited to the kinase domain of ROS1.

### Structural Modeling and Molecular Dynamic Simulation Studies

The ROS1 kinase domain was modeled in the DFG-in and DFG-out conformations using YASARA Version 20.12.24 as we described previously in Somwar *et al (Somwar et al, 2020)*. Point mutations were introduced, molecular dynamics simulations performed, and principal component analysis conducted using ProDy (Bakan *et al*, 2011) (Bakan *et al*, 2014) with methods described in detail in our previous publication (Keddy *et al*, 2022).

### Global Proteomics and Phosphoproteomics

#### Disulfide bond reduction/alkylation

Protein solutions (200 μg) were in 50 mM HEPES at 2 μg/μl in 1.5 mL Eppendorf low-bind tubes. Disulfide bonds within the proteins were reduced by adding tris (2-carboxyethyl) phosphine to a final concentration of 5 mM and mixing at room temperature for 15 min. The reduced proteins were alkylated by adding 2-chloroacetamide to a final concentration of 10 mm and mixing in the dark at room temperature for 30 min. Excess 2- chloroacetamide was quenched by adding dithiothreitol to a final concentration of 10 mM and mixing at room temperature for 15 min.

#### Methanol/Chloroform precipitation and protease digestion

Alkylated samples were subjected to protein precipitation as follows: 400 μL of methanol was added to the sample and vortexed for 5, 100 μL of chloroform was added to the sample and vortexed for 5 s, 300 μL of water was added to the sample and vortexed for 5 s, and the samples were centrifuged for 1 min at 14,000 g. The aqueous and organic phases were removed, leaving a protein wafer in the tube. The protein wafers were washed with 400 μL of methanol and centrifuged at 21,000 g at room temperature for 2 min. The supernatants were removed, and the pellets were allowed to air dry but not to complete dryness. The samples were resuspended in 70 μL 100 mM HEPES (pH 8.5) and digested with rLys-C protease (100:1, protein to protease) with mixing at 37 ℃ for 4 hr. Trypsin protease (100:1, protein to protease) was added and the reaction was mixed overnight at 37 ℃.

#### TMTpro15plex labeling

TMTpro16plex labeling reagent (Pierce, 500 μg) was brought up in 30 μL acetonitrile and added to the digested peptide solution (200 μg) yielding a final organic concentration of 30% (v/v) and mixed at room temperature for 1 hr. A 2 μg aliquot from each sample was combined, dried to remove the acetonitrile, processed with a C18 ZipTip (Millipore) and analyzed via LC/MS as a “label check”. Equalization ratios and labeling efficiency were determined and the reactions were quenched with the addition of hydroxylamine to a final concentration of 0.3% (v/v) for 15 min with mixing. The TMTpro labeled samples were pooled at 1:1 ratio based on the equalization ratios from the label check and concentrated in a speedvac to remove acetonitrile. The material was desalted with a SepPak C18 3mL cartridge (Waters) with the elution being split into two fractions, with 1 mg being used for pTyr enrichment and 2 mg being fractionated by basic reverse phase fractionation. Both fractions were taken to dryness using a SpeedVac.

#### bRP Fractionation

The 2 mg combined TMT sample was resuspended in 100 µL 10 mM ammonium bicarbonate pH 8. The material was loaded on to a Zorbax 2.1 x 150 mm (5 µm particle size) Extend-C18 column (Agilent) for basic reverse-phase fractionation. The sample was gradient-eluted from the column at a flowrate of 250 µL/min over 55 minutes using a combination of solvents “A” (10 mM ammonium carbonate) and “B” (acetonitrile). The gradient used was as follows: from 0 to 5 minutes, “B” was held at 1%, from 5 to 55 minutes, “B” varied from 5% to 40%, followed by an increase to 90% “B” over 5 minutes, and then a hold for an additional 5 minutes at 90% “B”. The UV signal was monitored at 210 nm. Ninety-six 50 second fractions were collected and combined into 24 pools by concatenation (where every 24^th^ fraction was combined into a pool). The pools were subsequently taken to near-dryness by vacuum centrifugation. Pools were brought up to 100 µL in 98/2/0.1% (v:v) water:acetonitrile/formic acid and 5% from each pool was analyzed by LC-MS with an Orbitrap Eclipse. The pools were further combined to 12 fractions and taken to dryness by vacuum centrifugation for subsequent IMAC enrichment

#### IMAC Enrichment

A 100 µL slurry of 5% Ni-NTA magnetic beads (Qiagen, part# 36113) was placed in 12 1.5 mL Eppendorf tubes and the tubes were placed in a magnetic stand and the supernatant was removed. The beads were rinsed three times with 100 µL water, each time placing the samples in the magnetic stand and removing the supernatant. A 100 µL solution of 40 mM EDTA was added to the beads, vortexed, mixed at ∼ 1400 rpm for 30 minutes at room temperature, and the supernatant was removed. The beads were rinsed three times with 100 µL water, each time placing the sample on the magnetic stand and removing the supernatant. A 100 µL solution of 10 mM FeCl3 was added to the beads, vortexed and mixed at ∼ 1400 rpm for 30 minutes at room temperature, and the supernatant was removed. The beads were rinsed three times with 100 µL water, each time placing the sample on the magnet and removing the supernatant. The beads were rinsed three times with 100 µL 80% ACN/0.1% TFA before resuspending them in 100 µL 80% ACN/0.1% TFA. The bRP fractionated TMT labeled peptides (12 fractions) were resuspended in 200 µL 80% ACN/0.1% TFA and added to the resuspended beads. The samples were vortexed and mixed at ∼ 1400 rpm for 30 minutes at room temperature. Samples were spun down quickly at 1000g for 10 s and placed on the magnetic stand to remove the supernatant. The samples were washed three times with 300 µL 80% ACN/0.1% TFA, each time placing the samples on the magnetic stand and removing the supernatant. Phosphopeptide elution was carried out by adding 200 µL of 70% ACN/1% ammonium hydroxide to the beads, mixing at room temperature for 1 min, placing the samples on the magnetic stand, and transferring the supernatants into fresh Eppendorf tubes containing 60 µL of 10% formic acid and mixing. The samples were taken to dryness by vacuum centrifugation.

#### Phosphotyrosine peptide enrichment

The 1 mg TMT-labeled peptide fraction underwent phosphotyrosine peptide enrichment using the PTMscan HS Phospho-Tyrosine (P-Tyr-1000) Kit (Cell Signaling Technology) following the manufacturer’s instructions. Eluted peptides were desalted on an Ultra-micro Spin Column (Harvard Apparatus) and taken to dryness by vacuum centrifugation prior to mass spectrometry analysis. *LC/MS*: The generated basic reverse phase fractions were brought up in 2% acetonitrile in 0.1% formic acid (20 μL) and analyzed (2 μL, 18 μL IMAC and pY) by LC/ESI MS/MS with a Thermo Scientific Easy1200 nLC (Thermo Scientific, Waltham, MA) coupled to a tribrid Orbitrap Eclipse with FAIMS pro (Thermo Scientific, Waltham, MA) mass spectrometer. In-line de-salting was accomplished using a reversed-phase trap column (100 μm × 20 mm) packed with Magic C18AQ (5-μm 200Å resin; Michrom Bioresources, Auburn, CA) followed by peptide separations on a reversed-phase column (75 μm × 270 mm) packed with ReproSil-Pur C18AQ (3-μm 120Å resin; Dr. Maisch, Baden-Würtemburg, Germany) directly mounted on the electrospray ion source. A 120-minute gradient from 4% to 44% B (80% acetonitrile in 0.1% formic acid/water) at a flow rate of 300 nL/minute was used for chromatographic separations. A spray voltage of 2300 V was applied to the electrospray tip in-line with a FAIMS pro source using varied compensation voltage -40, -60, -80 while the Orbitrap Eclipse instrument was operated in the data-dependent mode, MS survey scans were in the Orbitrap (Normalized AGC target value 300%, resolution 120,000, and max injection time 50 ms) with a 3 sec cycle time and MS/MS spectra acquisition were detected in the Orbitrap (Normalized AGC target value of 250%, resolution 50,000 and max injection time 100 ms) using higher energy collision-induced dissociation (HCD) activation with HCD collision energy of 38%.

### Data analysis

Data analysis was performed using Proteome Discoverer 2.5 (Thermo Scientific, San Jose, CA). The data were searched against a Mouse database (UP00000598 Human 030721) that included common contaminants (CRAPome) (Mellacheruvu *et al*, 2013). Searches were performed with settings for the proteolytic enzyme trypsin. Maximum missed cleavages were set to 2. The precursor ion tolerance was set to 10 ppm and the fragment ion tolerance was set to 0.6 Da. Dynamic peptide modifications included oxidation (+15.995 Da on M). Dynamic modifications on the protein terminus included acetyl (+42.-11 Da on N-terminus), Met-loss (-131.040 Da on M) and Met-loss+Acetyl (-89.030 Da on M) and static modifications TMTpro (+304.207 Da on any N-termius), TMTpro (+304.207 DA on K) and carbamidomethyl (+57.021 on C). IMAC searches included phosphorylation (+79.966 Da on S, T, and Y) as a dynamic modification. Sequest HT was used for database searching. IMP-ptmRS was used for phospho searches. All search results were run through Percolator for scoring. Finally, downstream analysis using the KSEA and Causalpath software packages were run according to previously published protocols (Babur *et al*, 2021; Casado *et al*, 2013; Luna *et al*, 2021; Wiredja *et al*, 2017).

### *In vivo* efficacy studies

All animal model studies were conducted in accordance with the Animal Welfare Act (AWA), Public Health Service (PHS), the United States Department of Agriculture (USDA), and under auspices of an approved protocol from the OHSU Institutional Animal Care and Use Committee (IACUC). Four-to eight-week-old female and male athymic nude (Nu/j) mice were obtained from The Jackson Laboratory (Bar Harbor, ME) and housed and handled under specific pathogen-free conditions in the University’s Animal Care Facilities. After an initial two-week environmental adjustment period, mice were placed under anesthesia using 2% isoflurane/oxygen, weighed, and ear punched for identification purposes. Tumor cells (1 – 5 x 10^6^ were mixed with 50 μl of matrigel and injected subcutaneously into the left or right flank. Animals were allowed to recover under supervision before being returned to animal facilities. Injected animals were checked daily until tumor were palpable nodules, at which time both animal weight and tumor size were measured thrice weekly using balance and a digital caliper (cat 14-648-17, Fisher Scientific, Federal Way, WA). Tumors were allowed to grow until they reached the humane limit of 1500- 2000 mm^3^ at which time animals were sacrificed and tumors were collected. At collection, tumors were washed in PBS and dissected with sterile scalpel blades and processed as follows: half the tumor was fixed in 10% normal formalin (24 hours then placed in 70% EtOH) for immunohistochemistry, the rest was divided into aliquots for freezing in liquid nitrogen or used for tumor-derived cell line generation (D2113N-P1). Initial studies of tumor formation were carried out in 4 female Nu/j mice (JAX lab) 4 to 6 weeks of age and repeated in male mice as shown in Supplementary Data.

Once tumor formation was documented, a follow-up study to test effect of TKI inhibitors on tumor growth was performed. ROS1^D2113N^ mutated NIH 3T3 cells were transduced with Firefly Luciferase lentiviral particles and selected with blasticidin (10 µg/ml) according to the manufacturer protocol (Cellomics Technology, Rockville, MD). Using luciferase-enabled monitoring of tumor growth using non-invasive bioluminescent imaging with the IVIS® Spectrum *in vivo* imaging system (Caliper LifeSciences, Hopkinton, MA). For this, luciferin dissolved in PBS was administered to mice by IP injection at a final dose of 150 mg/kg. After 6 minutes anesthesia was induced by 2.5% isoflurane/2.5% oxygen and mice were imaged with data being recorded. For the TKI treatments, 26 Nu/j mice (13 females/13 males) were injected with luciferase expressing ROS1-D2113N mutated NIH 3T3 cells (1 x 10^6^) in the right flank after animals had been anesthetized with 2.5% isoflurane/2.5% oxygen. *In vivo* tumor measurement began 3 days after injection and caliper measurements were initiated as soon as palpable nodules were noted. TKI treatment began 29 days after initial injection when tumor volume was reliably ∼90-120 mm^3^). At this time, animals were randomly assigned to TKI treatment groups after photon measurements were ranked from high to low to ensure higher expressing tumors based on photon emission were randomly dispersed amongst lower expressing tumors. Control animals were treated with vehicle by gavage (ethanol/PEG200/Water 10%/40%/50%). The lorlatinib treatment group animals was 3mg/kg lorlatinib daily via oral gavage (lorlatinib formulation was in ethanol/PEG200/Water 10/40/50). Crizotinib (100mg/kg) treatment was also via oral gavage; crizotinib was formulated in 0.5% methyl cellulose/0.5% Tween-80. Mice were monitored for tumor growth by caliper measurements twice weekly and once a week assessed with the IVIS® Spectrum *in vivo* imaging system, until tumor volume approaches 2000 mm^3^, the humane end point of the study.

### Statistical analysis

Mean ± SEM are shown unless otherwise stated. Student’s t-test, one-way, or two-way ANOVA with multiple comparisons tests were used and specified in the figure legends. P values < 0.05 were deemed statistically significant. Asterisks used in figures correspond with descriptions in figure legends that denote level of statistical significance. For global proteomics and phosphoproteomics, both pairwise two-sample *t* tests and Wilcoxon rank-sum test were conducted (R package) to delineate differential expression proteins and Benjamini-Hochberg procedure was used for controlling the false discovery rate (FDR). All data were plotted and analyzed using GraphPad Prism v9.3 (RRID:SCR_002798).

## Results

### Functional screening of clinical tumor sequencing data identifies several ROS1 variants that exhibit increased catalytic activity relative to wildtype ROS1

Our primary goal was to identify potential gain-of-function mutations in the receptor tyrosine kinase, *ROS1*, that drive oncogenic transformation. We queried patient sample cohort in the AACR Genie database (genie.cbioportal.org), a publicly available portal that contains clinical tumor sequencing data from multiple cancer centers and cancer types (Cerami *et al*., 2012; Consortium, 2017; Gao *et al*., 2013). We found that 3.5% of samples (total number 138,915) in the AACR Genie database contained *ROS1* alterations (Supplementary Table 1) (Cerami *et al*., 2012; Consortium, 2017; Gao *et al*., 2013). We prioritized somatic mutations for laboratory testing based on three criteria. First, we included only *ROS1* tyrosine kinase domain (TKD; amino acids 1945-2222 of ROS1) nonsynonymous missense mutations confirmed to be somatic or with mean allele frequency (MAF) < 0.01, 2). We excluded ROS1 mutations from tumor samples that harbored >15 total alterations (>15 total mutation burden for samples where information was available). We used the algorithm ‘Sorting Intolerant From Tolerant’ (SIFT) that predicts potential impact of mutation on protein function and included ones that showed deleterious impact. This filtering process yielded 33 missense somatic mutations in the *ROS1* TKD that we functionally characterized. As shown in **Figure 1A**, these 33 mutations do not cluster in any subdomain within the ROS1 TKD, such as around the P-loop, C-helix, HRD motif, or the activation loop (A-loop).

**Figure 1:**
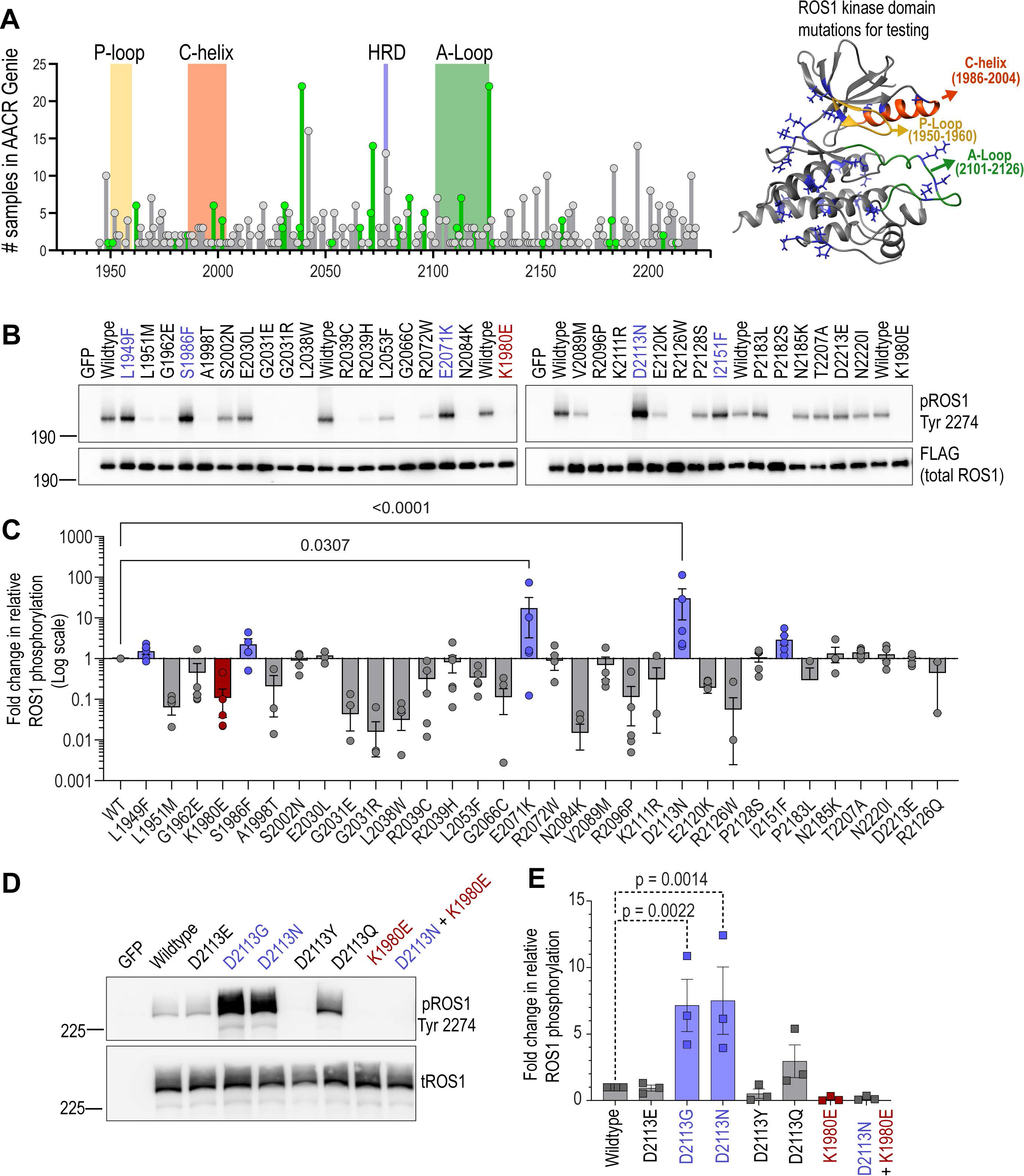
Functional screening of missense ROS1-TKD mutations identifies potential gain of function variants. **A.** Lollipop diagram shows missense mutations in the ROS1 TKD from AACR Genie. Y-axis represents number of samples with mutation at kinase domain position while X-axis shows amino acid position of ROS1 (Kinase Domain: AA 1945-2222). Grey colored mutations did not pass the filtering criteria while green colored mutations were included in functional testing. Key kinase domain motifs are shaded and labeled as indicated. **B.** Representative immunoblot of transfected missense ROS1 TKD mutations in transfected HEK- 293A lysates. **C.** Densitometry analysis of immunoblots (n=3) as represented in (B). Fold-change in relative ROS1 phosphorylation is calculated by first calculating the ratio of phospho-ROS1 to total ROS1 protein for each variant and then normalizing to ROS1^WT^. **D.** Representative immunoblot of ROS1 D2113 position substitutions (E, G, Y, and Q as indicated) in HEK-293A cells. E. Densitometry of immunoblots (n=3) as represented in (D). pROS1- phospho-ROS1, tROS1- total-ROS1. One-way ANOVA and Sidak’s test were used to control for multiple comparisons. Asterisks indicate statistical significance (** P < 0.01; * P < 0.05; ns- P > 0.05).

Using site-directed mutagenesis, we cloned the 33 *ROS1* variants to compare to wildtype *ROS1* (ROS1^WT^) and a negative control, ROS1^K1980E^, which represents a kinase dead variant with inability to coordinate ATP. To assess gain-of-function, we transiently transfected these ROS1 constructs into HEK293T/17 cells and immunoblotted the resulting lysates with a phospho- specific ROS1 antibody that detects the C-terminal ROS1 autophosphorylation (Y2274); this phospho-site serves as a surrogate measure for kinase catalytic activity in our studies (**Fig. 1B**). The engineered kinase dead variant, ROS1^K1980E^, did not have any detectable kinase activity, as expected. The ROS1 amino acid changes of potential interest for gain of function were: L1949F, S1986F, E2071K, D2113N, and I2151F (**Fig. 1C**). Among these, ROS1^D2113N^ exhibited highest activity with a median 4.8-fold increase in auto-phosphorylation (range 2.02-114.3) relative to ROS1^WT^. To further investigate the functional impact of perturbing the D2113 position that resides in the activation loop of the kinase, we generated other substitutions, D2113E, D2113G and D2113Y that were subsequently also discovered in the AACR Genie sequencing data (Supplementary Table). Additionally, we engineered D2113Q to understand the structural impact of substituting an uncharged polar residue at this position. As an additional negative control, we designed a compound mutation, ROS1^D2113N/K1980E^. Transient transfection and immunoblotting for ROS1 autophosphorylation showed that conservative substitution of D2113E does not alter kinase behavior, D2113Y substitution with a large hydrophobic residue introduction was deleterious for catalytic function, D2113Q was also modestly activating and importantly the small hydrophobic substitution with D2113G robustly increased catalytic activity to the same extent as the D2113N (**Fig. 1D, E**). As hypothesized, kinase dead variant ROS1^K1980E^ exhibited no catalytic activity and the compound mutation D2113N/K1980E phenocopied, confirming the phosphorylation increase at ROS1 Y2274 is due to increase in intrinsic catalytic activity. Thus, our functional screening approach yielded the following potential gain-of-function variants that warranted further functional validation: ROS1 L1949F, S1986F, E2071K, D2113G/N, T2207A, I2151F. The location of these residues is mapped on the structural model of ROS1 kinase domain (Supplementary Figure 1A).

### Cell-based transformation and proliferation assays reveal that ROS1^D2113N^ is a gain-of-function missense variant that promotes ROS1 TKI-sensitive oncogenic growth

To determine if the increase in ROS1 catalytic activity is sufficient to induce neoplastic transformation, we employed the routinely used NIH-3T3 soft agar colony formation assay (Arai *et al*., 2013; Borowicz *et al*, 2014; Davare *et al*, 2018; Davare *et al*., 2013). Conferring anchorage independent cell growth is one of the in vitro hallmarks of the tumorigenic potential of oncogenes. To this end, we stably transduced ROS1 L1949F, S1986F, E2071K, D2113N, I2151F and ROS1^WT^ as well as negative control, ROS1^K1980E^, in NIH-3T3 cells and determined extent of autophosphorylation of selected ROS1 mutants in this independent model system (**Fig. 2A**). The extent of increase in ROS1 activity is reproducible in NIH-3T3 cells, with ROS1^D2113N^ demonstrating the highest catalytic activation as illustrated by ROS1 Y2274 autophosphorylation (**Fig. 2A**). In colony formation experiments, only the NIH-3T3 ROS1^D2113N^ cells induced significant (p < 0.0001) colony formation relative to NIH-3T3 empty vector (EV) and ROS1^WT^ cells (**Fig. 2B**). We note that overexpression of wildtype ROS1 also increased colony formation potential compared to ROS1^K1980E^ kinase dead or EV controls, albeit this was not significant. ROS1-induced colony formation was blocked by concurrent ROS1-TKI treatment with a relatively selective ROS1 inhibitor, lorlatinib; thus, confirming the contribution of ROS1 tyrosine kinase catalytic activity to this phenotype (**Fig. 2B**). ROS1 fusions are known oncogenes in cancer and we wanted to compare the oncogenic potential of the known SLC34A2-ROS1 fusion (denoted SLC-ROS1) to ROS1^D2113N^ in this experimental model system. As shown in **Figure 2C**, SLC-ROS1 is a superior oncogene with nearly five-fold higher colony count compared to ROS1^D2113N^; albeit both harbor transformative capacity compared to kinase dead ROS1^K1980E^ (**Fig. 2C**). To test the pharmacological targetability of ROS1^D2113N^ with ROS1 TKI, we treated NIH-3T3 ROS1^D2113N^ cells with 10, 100 or 500 nM of crizotinib (FDA-approved), entrectinib (FDA-approved) and lorlatinib (phase III clinical trial) followed by assessment of ROS1 autophosphorylation and effects on downstream effector pathway modulation (Supplementary Figure S1B). Treatment with crizotinib, entrectinib or lorlatinib caused dose-dependent inhibition of ROS1 autophosphorylation at Y2274. These data suggest that ROS1^D2113N^ regulates activation of SHP2, AKT and STAT3 phosphorylation on canonical activation sites since both these were sensitive to ROS1 TKI and extent of their inhibition correlated to decrease in ROS1 phosphorylation. In contrast, activation of Src, ERK1/2 (MAPK), and ribosomal protein S6 is not linked to ROS1^D2113N^ catalytic activity as they are insensitive to ROS1 TKI treatment (Supplementary Figure S1B). To preliminarily assess effect of ROS1 TKI crizotinib on cell growth in NIH-3T3 cells, we performed dose response cell viability assays with NIH-3T3 ROS1^D2113N^ cells compared to negative control, NIH-3T3 empty vector (EV) and three positive controls, CD74-ROS1 wildtype, CD74-ROS1 G2032R (ROS1 TKI resistant mutant), and SLC34A2-ROS1. While NIH-3T3 model is typically used for morphologic assessment, even in cell viability assays ROS1^D2113N^ cells exhibited comparable sensitivity to crizotinib as the fusion proteins. Notably, crizotinib response is on-target for ROS1 because the crizotinib resistant CD74-ROS1 G2032R fusion cell line had significantly reduced sensitivity as expected (Supplementary Figure S1C, D).

**Figure 2:**
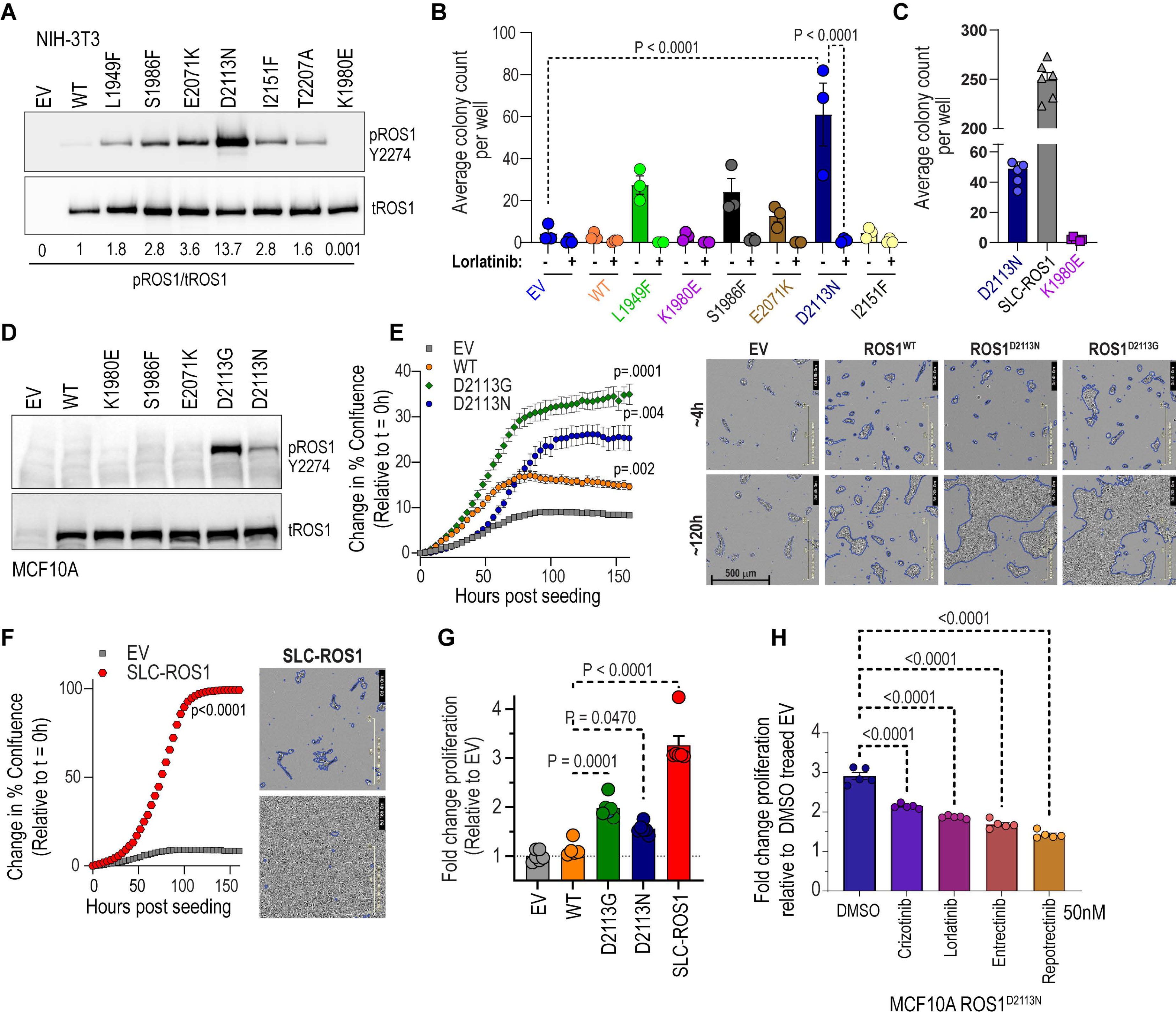
Cell-based transformation assays with catalytically activated ROS1 TKD variants. **A.** Immunoblot analysis of phospho-ROS1 and total ROS1 protein expression of NIH-3T3 cells stably transduced with empty vector (EV) or indicated ROS1 TKD mutants **B.** Soft-agar assay of NIH-3T3 ROS1 cell lines (n=3) treated with DMSO or 50 nM lorlatinib for 3 weeks. Two-way ANOVA with multiple comparisons test was used to determine statistical significance. **C.**Soft- agar assay of NIH-3T3 ROS1^D2113N^ compared with NIH-3T3 SLC-ROS1 fusion cells (positive control) or ROS1^K1980E^ (kinase-dead, negative control). **D.** Immunoblot analysis of phospho- ROS1 and total ROS1 protein expression in MCF10A cells stably transduced with indicated ROS1 variants. **E.**MCF10A proliferation assay with reduced EGF (0.5 ng/mL) as assessed via Incucyte live-cell imaging. Representative image taken from Incucyte platform with confluence mask definition (indicated by blue outline) was used to calculate % confluence values at indicated time point. Two-way ANOVA with Dunnett’s multiple comparisons used to test for statistical significance. **F.** MCF10A proliferation assay with reduced EGF (0.5 ng/mL) of cells transduced with SLC-ROS1 fusion relative to empty vector as assessed via Incucyte live-cell imaging. Representative image taken from Incucyte platform that demonstrates confluence mask definition used to calculate % confluence values at each timepoint. **G.** MCF10A stable cell line proliferation with reduced EGF (0.5 ng/mL) as assessed via CCK-8 colorimetric assay after one week. Absorbance measured at 490 nm. Values represent fold change relative to cells transduced with empty vector (EV). One-way ANOVA with Tukey’s multiple comparisons used to test statistical significance. **H.** MCF10A ROS1^D2113N^ proliferation with reduced EGF (0.5 ng/mL) media with or without DMSO (< 0.01%) or ROS1-TKIs treatment as indicated was measured with CCK-8 colorimetric assay after one week. Absorbance measured at 490 nm. Values represent fold-change relative to cells transduced with empty vector. One-way ANOVA with Tukey’s multiple comparisons used to test statistical significance. Asterisks indicate statistical significance (**** P < 0.001; ns- P > 0.05).

To validate transformative potential of ROS1^D2113N^ in a second independent model system, we utilized the immortalized MCF10A mammary epithelial cell line, which is dependent on epidermal growth factor (EGF) and insulin for proliferation. The MCF10A model is routinely used to characterize functional consequence and oncogenic potential of proto-oncogenes, including variants in PI3KCA (Debnath *et al*, 2003; Isakoff *et al*., 2005) and PDGFRA (Ip *et al*, 2018) recently. We generated stable MCF10A cells with viral transduction using constructs for EV and wildtype, K1980E, S1986F, E2071K, D2113N, D2113G. Intriguingly, in the stably transduced MCF10A cells, only ROS1^D2113N^ and ROS1^D2113G^ exhibited robust increase in Y2274 autophosphorylation compared to ROS1^WT^ (**Fig. 2D**). Based on confirmed increase in catalytic activity, ROS1^D2113N^ and ROS1^D2113G^ were selected to test in comparison with ROS1^WT^ and EV for assessing rate of cell proliferation under growth factor withdrawal conditions (See Methods). Using the Incucyte™ real-time cell-counting platform, we found that the rate of MCF10A proliferation was significantly increased with ROS1^D2113N^ and ROS1^D2113G^ expression as compared to ROS1^WT^ or EV expression (**Fig. 2E**). Representative images after 120 h of live imaging are shown in the right panel. Transduction of the known fusion oncogene, SLC-ROS1, into MCF10A cells significantly increased proliferation as expected (**Fig. 2F, G**), thus validating that the phenotype is linked to ROS1 catalytic activity. Supplementary Movies from live imaging studies over 164 h are presented in Supplementary Data. Finally, we tested a series of ROS1 TKIs (crizotinib, lorlatinib, entrectinib, repotrectinib, and cabozantinib) and found that each significantly attenuated MCF10A proliferation mediated by ROS1^D2113N^ (**Fig. 2H**). Taken together, our studies with both NIH-3T3 and MCF10A model systems point to ROS1^D2113N^ as a TKI-sensitive gain-of-function mutation that promotes increased oncogenic transformation and cell proliferation in different cell types.

### Structural modeling reveals that Asn substitution at the ROS1 D2113 position dramatically increasing local flexibility within that region of the A-loop in DFG-in kinase conformation

We aimed to understand the structural basis of the significant changes in ROS1 catalytic function induced by the D2113N mutation. To this end, we developed structural models of ROS1^WT^ and ROS1^D2113N^ in their ‘active’ or DFG-in and ‘inactive’ of DFG-out states. The highly conserved residues ‘DFG’ appear at the start of the activation loop and the positioning of the DFG motif broadly dictates the active or inactive conformation of the kinase (Modi & Dunbrack, 2019). Changes in catalytic activity due to mutations can result from altered structure, stability, and dynamics. Here, we performed molecular dynamics simulations of ROS1^WT^ and ROS1^D2113N^ DFG- in and DFG-out structural models to measure conformational changes and structural stability. Six representative conformations for four kinases (WT: DFG-in and DFG-out and D2113N: DFG-in and DFG-out) are depicted in Supplementary Figure S2A. Initial observations with root mean square deviation (RMSD) analysis revealed that the ROS1^D2113N^ mutation affords dramatically enhanced protein flexibility surrounding the activation loop (A-loop) region of the kinase, specifically in the DFG-in conformation (Supplementary Figure S2E), as compared to the P-loop or αC-helix subdomains in DFG-in (Supplementary Figure S2F, G)). Notably, ROS1^D2113N^ does not appear to influence RMSD in the DFG-out conformation for any sub-domains in the kinase domain (Supplementary Figure S2H-J). ROS1^WT^ and ROS1^D2113N^ remained stable during simulations with global changes in the kinase domain as shown in Total RMSD in Supplementary Fig. S2E, H (Total).

To quantify the differences in local residue fluctuation more closely, we examined the RMSF (Root Mean Square Fluctuation) at each residue over the simulation course: mobile residues would display higher averaged RMSF values while constrained ones would display lower RMSF values (Martínez, 2015). As shown in **Figure 3A**. RMSF analysis suggests that in ROS1^WT^ kinase domain, the DFG-in (active) and DFG-out (inactive) conformations display dramatic differences in activation loop mobility (black lines in Fig. 3A), with the DFG-out (inactive) conformation displaying far greater mobility of the activation loop than the DFG-in conformation. Intriguingly, ROS1^D2113N^ inverts this trend, where the DFG-in conformation’s A-loop displays larger flexibility. The P-loop and αC-helix, as seen in RMSD analysis, do not exhibit any substantial changes in fluctuations (**Fig. 3C, D**), strongly suggesting that the mutation-induced hypermobility is restricted to the A-loop. The lack of overt changes in the P-loop and αC-helix also serve as an internal negative control and a landmark for the kinase domain.

**Figure 3:**
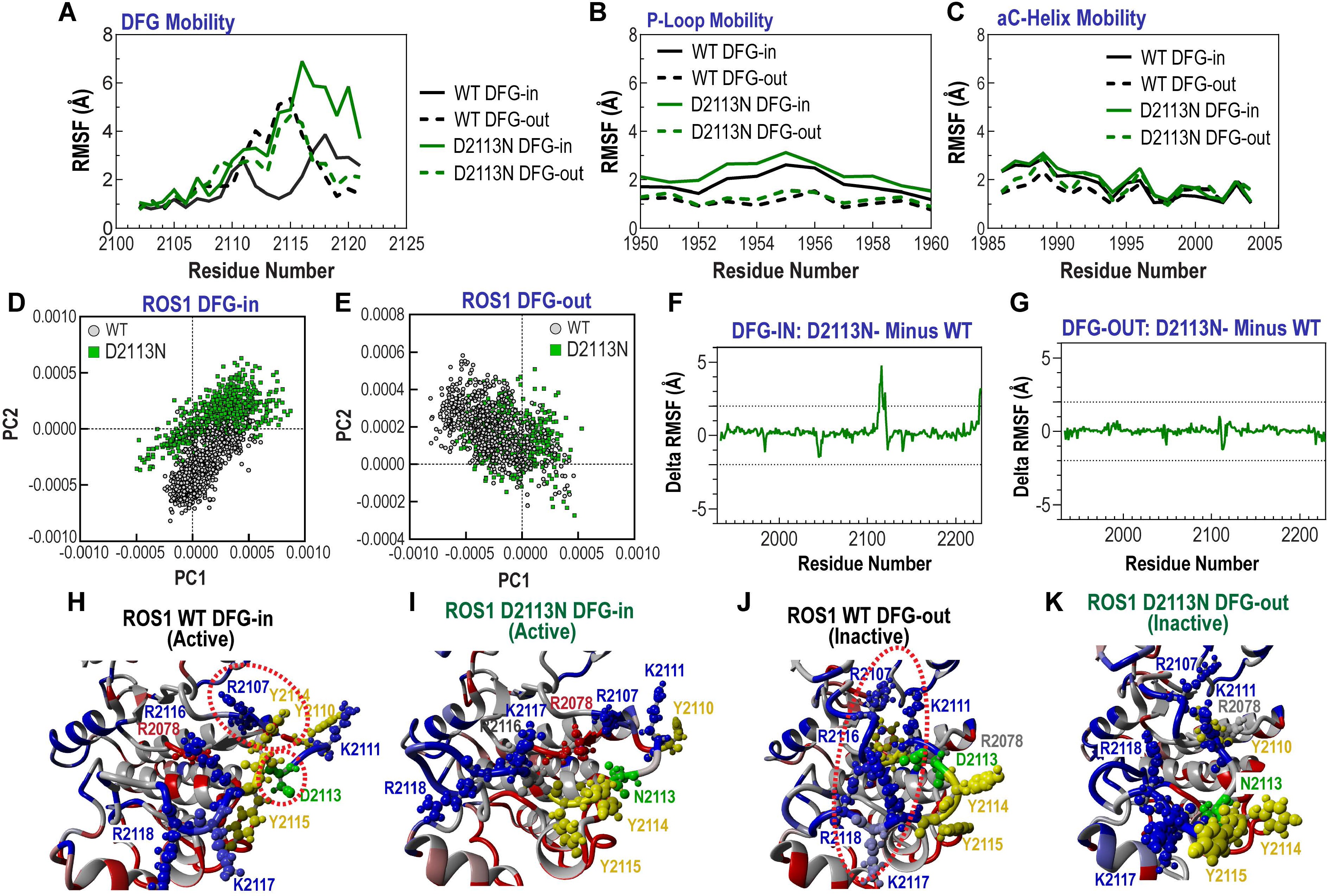
Structural modeling studies reveal dramatically increased dynamicity in the activation loop of ROS1^D2113N^ in DFG-in conformation. **A.** Root Mean Square Fluctuation (RMSF, Angstroms (Å) shown relative to residue location within A-loop from amino acid 2100 to 2125 in ROS1 wildtype (WT) and ROS1^D2113N^ (D2113N) DFG-in (active) and DFG-out (inactive) conformations. **B, C.**RMSF shown relative to residue location within P-loop from amino acid 1950-1960 (in B) and αC (aC) helix from amino acids 1985-2005 (in C) in ROS1 wildtype (WT) and ROS1^D2113N^ (D2113N) DFG-in (active) and DFG-out (inactive) conformations. **D, E.** Principal Component Analysis comparing relative distribution of ROS1 WT versus D2113N in DFG-in (active, panel D) and DFG-out (inactive, panel E) conformations. **F.** Difference in RMSF of D2113N – WT in DFG-in conformation and **G.** Difference in RMSF of D2113N – WT in DFG-in conformation. **H-K.** Structural model annotated and color-coded by residue for ROS1 WT DFG-in (H), ROS1 D2113N DFG-in (I), ROS1 WT DFG-out (J), ROS1 D2113N DFG-out (K).

Next, we hypothesized that these dramatic fluctuations in the A-loop of the D211N mutant kinase will ultimately influence the kinase pose. To assess this we performed principal component analysis (PCA) on the ensemble of poses and used clustering algorithm to discern hypothesized conformational clusters. Data in **Figure 3D, E** demonstrate that ROS1^D2113N^ exhibited the greatest difference in conformation relative to ROS1^WT^ when in the DFG-in as compared to the DFG-out position. These data are consistent with RMSD and RMSF findings and strongly indicate that the ROS1^D2113N^ in DFG-in conformation is a different kinase than ROS1^WT^.

To quantify the difference in A-loop fluctuations between ROS1^WT^ and ROS1^D2113N^, we subtracted the RMSF values between DFG-in and DFG-out confirmations, and between mutations (D2113 or N2113), and plotted the differences (**Fig. 3F, G**). An impressive, nearly 4-fold difference local fluctuation at or surrounding the D2113 residue is noted between ROS1^WT^ and ROS1^D2113N^ in the DFG-in conformation but not in the DFG-out conformation as expected (**Fig. 3F, G**). The modelled structures provide another intriguing angle. In general, these structures are colored by their surface electrostatic potential: blue suggests positive charge; red suggests negative charge; grey suggests neutral (**Fig. 3H-K**). The mutated residue (2113) is colored green, while phosphorylatable tyrosines (2110, 2114-5) are colored yellow. In the ROS1^WT^ DFG-out structure, the negatively charged D2113 may be pulled into a positively charged pocket produced by several basic residues (R2116, R2118, K2111, and possibly R2107) (**Fig. 3I**). This conformation appears to be stabilized not by interactions of the aspartate but by several cation:pi interactions with Y2114 (**Fig. 3H**). The ROS1^D2113N^ mutation alters the DFG-out inactive form moderately; here, the N2113 residue does not appear to come up as far into the pocket as its D2113 counterpart (**Fig. 3K**). However, this mutation dramatically alters the active DFG-in conformation; the model suggests that the cation pi interaction is not formed in the mutant kinase, and the activation loop autophosphorylation tyrosine, Y2114/Y2115 are in a different and potentially more exposed position relative to the kinase domain (**Fig. 3J**). Taken together, these structural modeling data suggest that the ROS1^D2113N^ mutation exerts a major dynamic effect by increasing the flexibility of the A-loop in the active, DFG-in, conformation.

### Phosphoproteomics reveal that ROS1^D2113N^ promotes increased PTPN11, GAB1, STAT, mTOR and JNK/SAPK activity relative to ROS1^WT^

With the discovery that ROS1^D2113N^ can promote oncogenic transformation in NIH-3T3 cells and increased cell proliferation in MCF10A cells, we wanted to study how ROS1^D2113N^ affects downstream signaling. To achieve this, we performed global proteomics and phosphoproteomics (IMAC and phosphotyrosine enrichment) on HEK-293A cells that were stably transduced with either ROS1^WT^ or ROS1^D2113N^ as described in Supplementary Fig. 3A. Both IMAC and pTyr phosphoproteomics revealed numerous statistically significant differences in a host of key downstream signaling proteins (**Fig. 4A-B**, Supplementary Fig. S4A). Increased phosphorylation and activation were observed in the RTK adapter proteins PTPN11 (SHP2) and GAB1, the transcription factors STAT1 and STAT3, RICTOR (contributor to the mTORC2 signaling complex), and ribosomal protein s6 (downstream effector of mTOR signaling) (**Fig. 4A-B**, Supplementary Fig. S4A).

**Figure 4:**
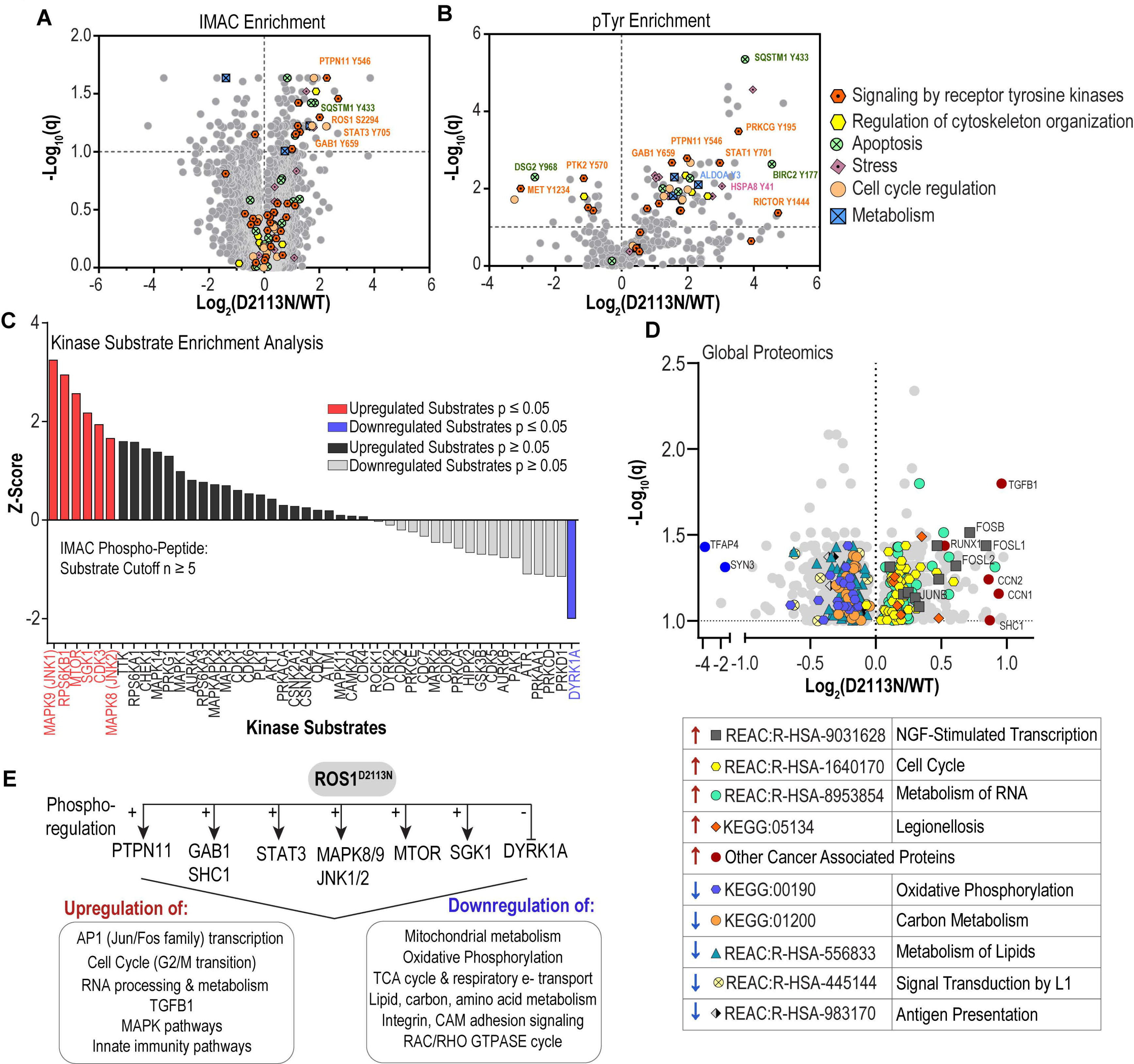
Phospho- and global proteomics identifies PTPN11, STAT, AP-1 transcription, and TGFB1 as ROS1^D2113N^-upregulated signaling effector pathways. **A-B.** Volcano plots of IMAC-enriched **(A)** and phospho-tyrosine-enriched **(B)** phosphoproteomic spectral counts from HEK-293A cells that express either ROS1^WT^ or ROS1^D2113N^. Selected phosphosites linked to certain cellular functions are annotated as indicated in graph legend. X-axis features log2- transformed ratio of normalized intensities of ROS1^D2113N^ relative to ROS1^WT^. The Y-axis features corresponding log10 (q) values with a false discovery rate of 10%. **C.** Kinase substrate enrichment analysis (KSEA) of IMAC phosphoproteomic data from **(A)** comparing ROS1^D2113N^ to ROS1^WT^. Red color indicates p ≤ 0.05 while positive z-score indicates that given kinase pathway is upregulated in ROS1^D2113N^ relative to ROS1^WT^. **D.** Volcano plots of global proteomics of HEK-293A cells that express either ROS1^WT^ or ROS1^D2113N^ with annotation of selected proteins linked to certain KEGG or Reactome-curated signaling pathways. Up arrows (red) signify upregulation while down arrows (blue) signify downregulation. The X-axis depicts difference between ROS1^D2113N^ and ROS1^WT^ using log2 transformed normalized intensities. The Y-axis shows corresponding log10 (q) values with a false discovery rate of 10%. **E.** Illustrated scheme broadly captures the significantly upregulated and downregulated effector proteins and/or signaling pathways (including KEGG and Reactome outputs) found in ROS1^D2113N^ cells relative to ROS1^WT^ cells. Raw proteomics counts are in Supplementary Tables. Data in panels A, B and D were analyzed and graphed using GraphPad Prism software.

To more broadly understand the effects of ROS1^D2113N^ on downstream signaling pathways, we performed Kinase Substrate Enrichment Analysis (KSEA) (Casado *et al*., 2013; Wiredja *et al*., 2017) on the phosphoproteomic data and found significant increase in phosphorylation of substrates of ribosomal S6 kinase, mTOR, SGK1, and JNK1/2 in ROS1^D2113N^ relative to ROS1^WT^ (**Fig. 4A- C**, Supplementary Fig. S4A). Global proteomics also revealed novel insight into pathways regulated by activated ROS1 receptor (**Fig. 4D**). Notable were increases in the AP-1 complex transcriptional factors, Jun/Fos, proteins involved with cell cycle progression, RNA metabolism, and upregulation of TGFB1 as well as Cysteine-rich protein 61 (CCN) family members, CCN1 and CCN2 (**Fig 4D)**. Intriguingly, proteins involved in oxidative phosphorylation and metabolism of carbon and lipids, as well as antigen presentation were downregulated. We also performed unbiased pathway analysis of the phosphoproteomics and global proteomics using Causalpath (Babur *et al*., 2021; Luna *et al*., 2021), which validated a link between ROS1^D2113N^ and increased SHP2, STAT, and TGFB1 signaling (Supplementary Fig. S3B). Intriguingly, minimal evidence was observed of any increased activity of either the canonical RAS-RAF-MEK-ERK or PI3K-AKT signaling pathways, which contrasts with the current thinking of signaling effects of ROS1 over activation as studied with ROS1 fusion oncogenes. Hence, our current working model of ROS1^D2113N^- mediated downstream signaling is summarized in **Figure 4E**, where increased activation of ROS1 provided by the D2113N mutant promotes increased signaling through STAT activation, mTOR signaling (likely mTORC2 with the presence of RICTOR phosphorylation), AP1 (Jun/Fos family) transcription, and TGFB1, with downregulation of non-glycolytic, mitochondrial metabolism.

### In vivo studies validate that ROS1^D2113N^ promotes tumor formation that is sensitive to inhibition by ROS1-TKI treatment

Finally, we assessed the tumorigenic potential conferred by the gain-of-function ROS1 mutations as well as *in vivo* efficacy of ROS1-TKI against this mutation. We implanted NIH-3T3 Empty Vector (EV), ROS1^WT^, ROS1^E2071K^, ROS1^D2113N^, ROS1^K1980E^ (negative control), and SLC- ROS1 (positive control) cells subcutaneously into Nu/J mice. The mice injected with NIH-3T3 cells that expressed the SLC-ROS1 fusion oncoprotein had the fastest growing tumors, which is expected given this is a known oncogene; similarly as expected, mice injected with either EV or ROS1^K1980E^ cells did not form tumors throughout the duration of the study (74 days) (**Fig. 5A-B**). ROS1^D2113N^ cells formed palpable tumors by day 32 and humane tumor volume limits were reached on day 52 (**Fig. 5A-B**). Intriguingly, mice injected with ROS1^WT^ or ROS1^E2071K^ cells also formed tumors albeit with greater latency (**Fig. 5A-B**). We examined expression of ROS1 and known phosphorylated effectors, pSHP2 and pSTAT3 in ROS1^D2113N^ tumors via immunohistochemistry (IHC) (**Fig. 5C**). The D4D6 ROS1 antibody was validated for its sensitivity and specificity for ROS1 using HEK293A cell pellets from stable cell lines expressing ROS1^WT^, CD74-ROS1 fusion, a truncated ROS1, ROS1^N2224*^ lacking the epitope for D4D6, and ETV6-NTRK3 fusion (Supplementary Fig. S5A). ROS1^D2113N^ tumors express ROS1 and have upregulated phosphorylation of SHP2 and STAT3, in line with phospho-immunoblot and proteomics findings (**Fig. 5C**). These results confirm that ROS1^D2113N^ is tumorigenic *in vivo*.

**Figure 5:**
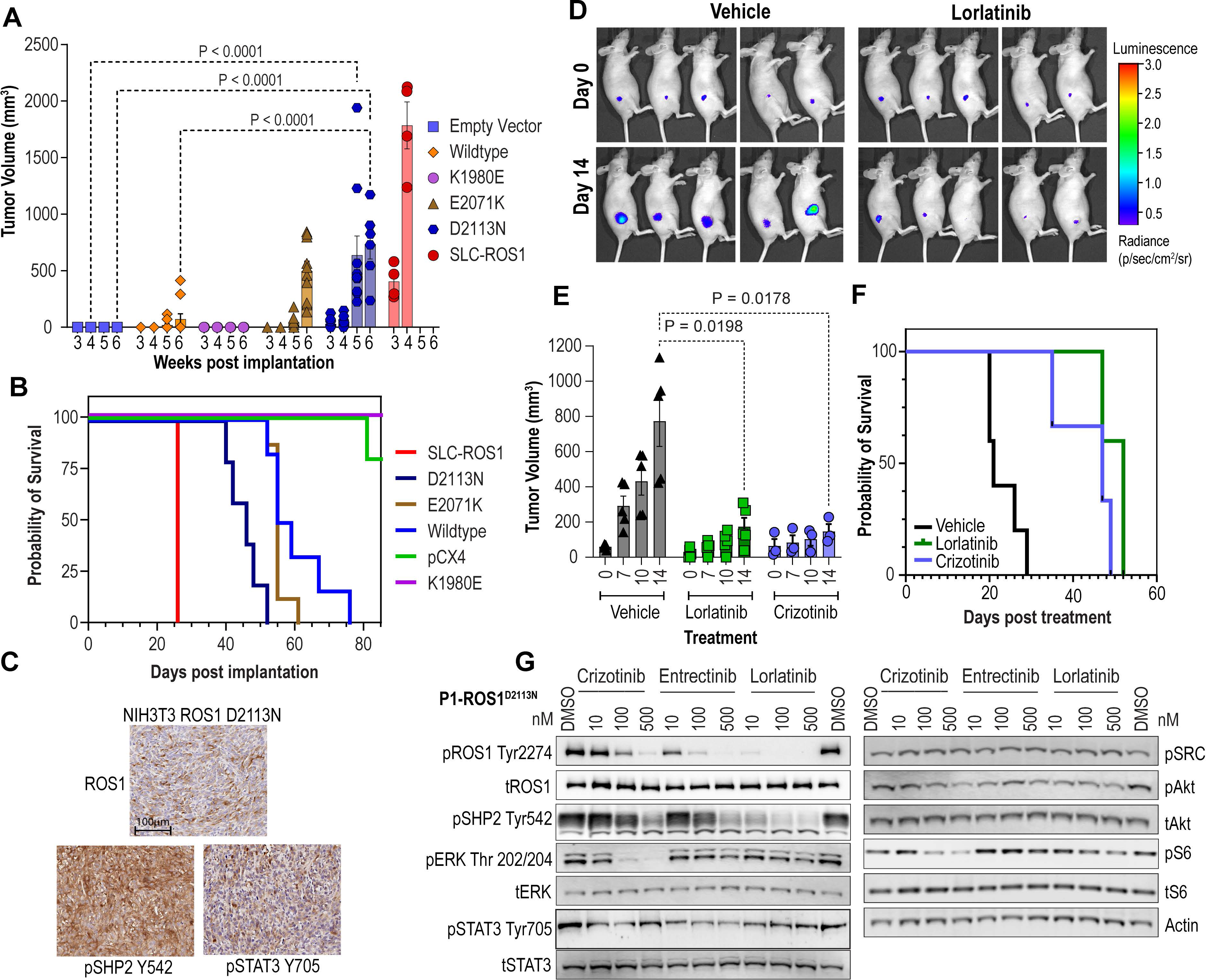
*In vivo* tumorigenesis studies confirm ROS1^D2113N^ as a gain-of-function oncogenic variant. **A.** Tumor volume (mm^3^) of NIH-3T3 empty vector, ROS1^WT^, ROS1^K1980E^, ROS1^E2071K^, ROS1^D2113N^, and SLC-ROS1 cells subcutaneously injected into flank of female Nu/J mice as monitored for 6 weeks. n=5 for all except SLC-ROS1 (n=4). **B.** Kaplan-Meier survival curve of mice described in (A). p < 0.0001. **C.** Representative images of immunohistochemistry of NIH- 3T3 ROS1^D2113N^ tumors probed for total ROS1, phospho-SHP2 (Y542), and phospho-STAT3 (Y705) expression. **D.** Representative images of photonic flux from luciferase-engineered ROS1^D2113N^ cell implanted into Nu/J mice and treated with Vehicle or lorlatinib (3 mg/kg) on days 0 and 14 of treatment. Non-invasive bioluminescent imaging achieved via IVIS Spectrum™ imaging platform. **E.** Tumor volume of NIH-3T3 ROS1^D2113N^ cells subcutaneously injected into female Nu/J mice (n = 3) and treated for 14 days with vehicle, crizotinib (100 mg/kg), or lorlatinib (3 mg/kg). **F.** Kaplan-Meier survival curve of mice described in (C). p < 0.0001. Asterisks indicate statistical significance (**** P < 0.001; * P < 0.05; ns- P > 0.05). **G.** Immunoblot analysis of the phosphorylated (p) and total (t) proteins from NIH-3T3 ROS1^D2113N^ P1 cell lysates prepared from cells treated with vehicle (DMSO) or 10, 100, or 500 nM of crizotinib, entrectinib, or lorlatinib for 4 hours.

We tested if pharmacological treatment with ROS1 TKI could attenuate the tumor growth driven by ROS1^D2113N^. For these studies, we expressed firefly luciferase via lentiviral transduction into NIH-3T3 ROS1^D2113N^ cells and performed flank injections in female Nu/J mice for tumor formation. Once palpable tumors were 90-120 mm^3^ as measured with caliper, we randomized mice into vehicle, crizotinib (100mg/kg, po, qd), and lorlatinib (3mg/kg, po, bd) treatment groups (n=3) (**Fig. 5D**). Over two weeks of treatment, both crizotinib and lorlatinib significantly attenuated ROS1^D2113N^-driven tumor growth in mice relative to vehicle (**Fig. 5D-E**) and increased survival relative to vehicle treatment (**Fig. 5F**). To ensure that ROS1^D2113N^ is tumorigenic in both sexes and test TKI effects, we repeated the *in vivo* TKI efficacy study in male Nu/J mice (Supplementary Figure S5B). These data show that ROS1^D2113N^ is tumorigenic independent of sex (formed tumors in male mice) and crizotinib effectively attenuates tumor formation; in these studies with male mice, lorlatinib was effective at inhibiting tumor growth but did not achieve statistical significance. We created a new tumor-derived cell line from the NIH-3T3 ROS1^D2113N^ tumors called P1- ROS1^D2113N^ (Supplementary Figure 5D) and confirmed that TKI treatment had on-target activity against ROS1 and associated downstream signaling as seen in the pre-tumor NIH-3T3 ROS1^D2113N^ cells via immunoblotting (**Fig. 5G**). In alignment, dose response cell viability with crizotinib also showed that P1-ROS1^D2113N^ cells derived from tumors retained dependence on ROS1 signaling (Supplementary Fig. S5E). Therefore, our data suggests that ROS1^D2113N^ drives tumor formation that can be attenuated or blocked with ROS1-TKI, *in vivo*, supporting the data from cell-based *in vitro* models.

## Discussion

Receptor tyrosine kinase (RTK) domain organization consists of an extracellular ligand- binding region (ECD), a single pass transmembrane helix, and an intracellular portion that features a juxtamembrane domain (JM), tyrosine kinase domain (TKD) and a C-terminal domain (CTD). RTKs normally serve to link extracellular signals to intracellular signaling pathways that affect cell proliferation, differentiation, and changes in metabolism (Schlessinger, 2014). It is now well established that aberrant constitutive RTKs catalytic activity upregulates downstream signaling pathways to promote uncontrolled cell proliferation and survival (Du & Lovly, 2018; Fleuren *et al*, 2016; Sanchez-Vega *et al*, 2018), two key hallmarks of cancer (Hanahan & Weinberg, 2011). Multiple mechanisms exist by which RTKs become hyper-activated. First, many RTK genes are involved in chromosomal rearrangements that are transcribed and translated into chimeric fusion oncoproteins that feature constitutively active TKDs which drive tumor formation in a wide variety of liquid and solid tumors (Rikova *et al*, 2007). Various groups have found a number of 5’ partner genes that can fuse via translocation or intrachromosomal deletion to the intracellular regions of RTK genes; in particular, studies have identified that *ROS1* can form fusion oncogenes with more than 55 partner genes (Drilon *et al*., 2021). A second mechanism of RTK overactivation involves either complete or partial RTK gene amplification that leads to increased RTK expression and downstream cell signaling. Notable examples include *HER2* amplification in breast cancer (Paik *et al*, 2008), *EGFR* amplification in gliomas (Ni *et al*, 2022; Smith *et al*, 2001), and *MET* amplification in non-small-cell lung cancer (NSCLC) (Drilon *et al*, 2017a). Finally, gain-of-function, nonsynonymous point mutations are also established mechanisms leading to RTK overactivation in cancer. While gain-of-function RTK mutations have been functionally characterized in both the ECD and intracellular regions of several RTKs, including EGFR, MET, RET and ALK, many mutations detected via clinical sequencing often group in the TKD, specifically in areas adjacent to the ATP-binding pocket and activation loop in these RTKs (Bresler *et al*, 2014; Lahiry *et al*, 2010; Medves & Demoulin, 2012).

A notable bottleneck limiting the translation of clinical cancer genomic sequencing data is the lack of functional classification of variants of unknown significance as druggable oncogenes or passengers. Thus, validating putative gain-of-function mutations represents an important effort for maximizing the clinical utility of NGS data. To this end, multiple *in silico* algorithms were developed that predict the effect of mutations on protein structure and function (Reva *et al*., 2007, 2011; Sim *et al*., 2012; Thusberg & Vihinen, 2009; Vaser *et al*., 2016); however, in our view, these algorithms are not yet exact enough to replace wet-lab functional validation. Indeed, risk of exclusive reliance on *in silico* approaches is revealed via this example: ALK F1174L, an oncogenic mutation in neuroblastoma (George *et al*, 2008; Mossé, 2016), is classified as a “neutral” functional impact variant by the popular Mutation Assessor algorithm. Thus, without laboratory-based validation, ALK F1174L would not be a biomarker for response to next-generation ALK TKI. *In silico* approaches may have a role in the future with improved functionality that includes machine learning or artificial intelligence paradigms that are rigorously trained on functionally validated datasets.

In this study, we discovered a gain-of-function missense mutation, ROS1^D2113N^ in the orphan RTK, *ROS1*; no activating mutations have previously been described for this RTK. Importantly, we observed that treatment with the ROS1-TKIs crizotinib (FDA-approved) and lorlatinib (phase III clinical trials) significantly attenuated ROS1^D2113N^ driven tumor growth, thus representing a potential therapeutic modality in tumors harboring this mutation. We found ROS1^D2113N^ has oncogenic effects in diverse cell types. ROS1^D2113N^ was detected in melanoma and oligodendroglioma in the AACR Genie database (Cerami *et al*., 2012; Consortium, 2017; Gao *et al*., 2013). This potentially suggests that activated ROS1 could promote cancer associated pathogenic pathways in a variety of cancer cell lineages, in concordance with the occurrence of ROS1 fusion oncogenes in highly diverse adult and pediatric cancers. More studies of ROS1^D2113N^ in various cell types would be beneficial to validate that TKI treatment against this variant can be beneficial in a diverse set of cancers. Broadly, these results establish that gain-of-function mutations in *ROS1* can drive cancer formation and that FDA-approved targeted therapy can potentially be leveraged in an expanded cohort of cancer patients.

The ROS1 receptor is poorly understood and accordingly signaling pathways activated by a constitutively active ROS1 RTK were unknown. Our proteomics and phosphoproteomics data for the first time permit a glimpse into the window of how the gain of function conferred by ROS1^D2113N^ affects cell-signaling pathways. A surprising finding for us was a lack of RAS-RAF- MEK-ERK and PI3K/AKT pathway activation in ROS1^D2113N^ cells as compared to ROS1 fusions where these pathways are established harbingers of ROS1 fusion mediated oncogenesis.

Intriguingly, we instead found that ROS1^D2113N^ predominantly promoted upregulation of JNK signaling and its downstream effectors the AP-1 transcription factor family consisting of Jun and Fos. Another unexpected finding, in comparison to ROS1 fusions is the upregulation of TGFBR ligand TGFB1, and the CCN family of proteins. Previous literature has demonstrated the link between activation of JNK family kinases and AP-1 transcriptional activity (Karin, 1995), of TGFβ upregulation by AP-1 transcription (Derynck *et al*, 2021), and notably, upregulation of CCN proteins by TGFβ (Nakerakanti *et al*, 2011; Tejera-Muñoz *et al*, 2021). Both TGFβ and the CCN family of proteins are linked to poorer prognosis in cancer through their effects on promoting stem cell-like behavior, decreased cell adhesion, and angiogenesis in diverse cancer types (Haque *et al*, 2011; Lau, 2012; Massagué, 2008). Further, this signaling axis is connected to increased cancer cell invasiveness and metastatic potential (Kim *et al*, 2018). This newfound link between activated ROS1 receptor and TGFβ has potential implications for therapy, as targeting of TGFβ signaling may provide an additional, potentially combinatorial avenue to target ROS1 mutation driven cancers. Furthermore, role of the full-length ROS1 RTK in normal physiology is poorly understood due to a dearth of studies devoted to the receptor in recent years. Indeed, the ligand for ROS1 is still unclear and our current understanding of ROS1 function relies solely on interrogation of ROS1 fusion proteins in cancer. Our data suggest that signaling patterns observed with ROS1 fusions do not necessarily recapitulate the signaling mediated by activation of the full- length ROS1. Hence, studies focusing on ROS1 signaling through alternative pathways such as TGFβ may yield greater insight into the physiological role of this orphan receptor.

Structural modeling gives initial insight into the potential mechanism via which ROS1^D2113N^ increases catalytic activity of ROS1 tyrosine kinase. Our data point to the dramatic impact of this mutation on local structural changes that lead to enhanced mobility of the region of A-loop that D2113 resides in. Notably, D2113 is adjacent to key auto-phosphorylation sites, Y2114 and Y2115 that are thought to be required for sustained catalytic activation and maintenance of the kinase structure in the DFG-in or open/active kinase conformation. Thus, we can theorize that these alterations in the A-loop surrounding D2113 residue may increase propensity of the tyrosine kinase to stay in the DFG-in like state resulting in presumed enhanced ATP binding and/or accessibility for both intrinsic (Y2114/Y2115) and extrinsic substrates such as effector proteins. We previously showed that ROS1^D2113N^ and to an even greater extent, ROS1^D2113G^, confers resistance to type II binding mode ROS1-TKI cabozantinib, in the context of ROS1 fusion protein, CD74-ROS1 (Davare *et al*., 2015). Briefly, type II inhibitors have higher affinity and binding preference for the DFG-out (inactive) kinase conformation. We established cabozantinib as a type II ROS1 inhibitor. In contrast, type I inhibitors have preferential binding to the DFG-in (active) kinase conformation, and thus are frequently referred to as ATP-competitive inhibitors. In this previous study aimed at profiling TKI resistance, the D2113N/G substitutions reduced or abrogated cabozantinib sensitivity of ROS1, which provides strong independent support for the conclusion that these D21113 position mutations force the kinase into a DFG-in or constitutively ‘active’ conformation that is not favorable for type II inhibitor binding. From a translational perspective, in the future it will be important to test if ROS1^D2113N^ in the full-length receptor context is also resistant to type II ROS1-TKI treatment akin to what we observed in fusion oncoprotein setting. Thus, more work in this area could greatly benefit future strategies to optimize TKI treatment for these patients. Additionally, future crystallographic or cryo-electron microscopy studies of mutant ROS1 kinase will be essential to confirm the findings from our modeling studies and offer higher resolution and deeper understanding of the impact of these mutations on ROS1 kinase structure-function relationship.

Ultimately, we hope that deconvolution of gain of function mutations amongst the plethora of reported variants of unknown significance in actionable genes such as ROS1 may enable an expanded cohort of patients to benefit from targeted kinase inhibitors to improve their outcomes.

## Acknowledgements

This project was supported with funding from the National Institutes of Health (NRSA: F30CA247253 (Iyer), NIH/NCI R01CA233495 (Davare) and RSG-19-082-01-TBG (Davare; American Cancer Society)) as well as the OHSU Doernbecher Children’s Hospital Foundation. We also would like to acknowledge the American Association for Cancer Research and its financial and material support in the development of the AACR Project GENIE registry, as well as members of the consortium for their commitment to data sharing. Interpretations are the responsibility of study authors. We would also like to acknowledge the Fred Hutchinson Cancer Center Proteomics & Metabolomics core service, including Phil Gafken, Chenwei Lin, and Lisa Jones for their support in performing the global proteomics and phosphoproteomics experiments.

## Notes

### Competing Interest Statement

The authors have declared no competing interest.

