## Supplementary Figures and Legends for "Discovery of oncogenic ROS1 missense mutations with sensitivity to tyrosine kinase inhibitors"

#### Supplementary Figures and Figure Legends

#### Figure S1

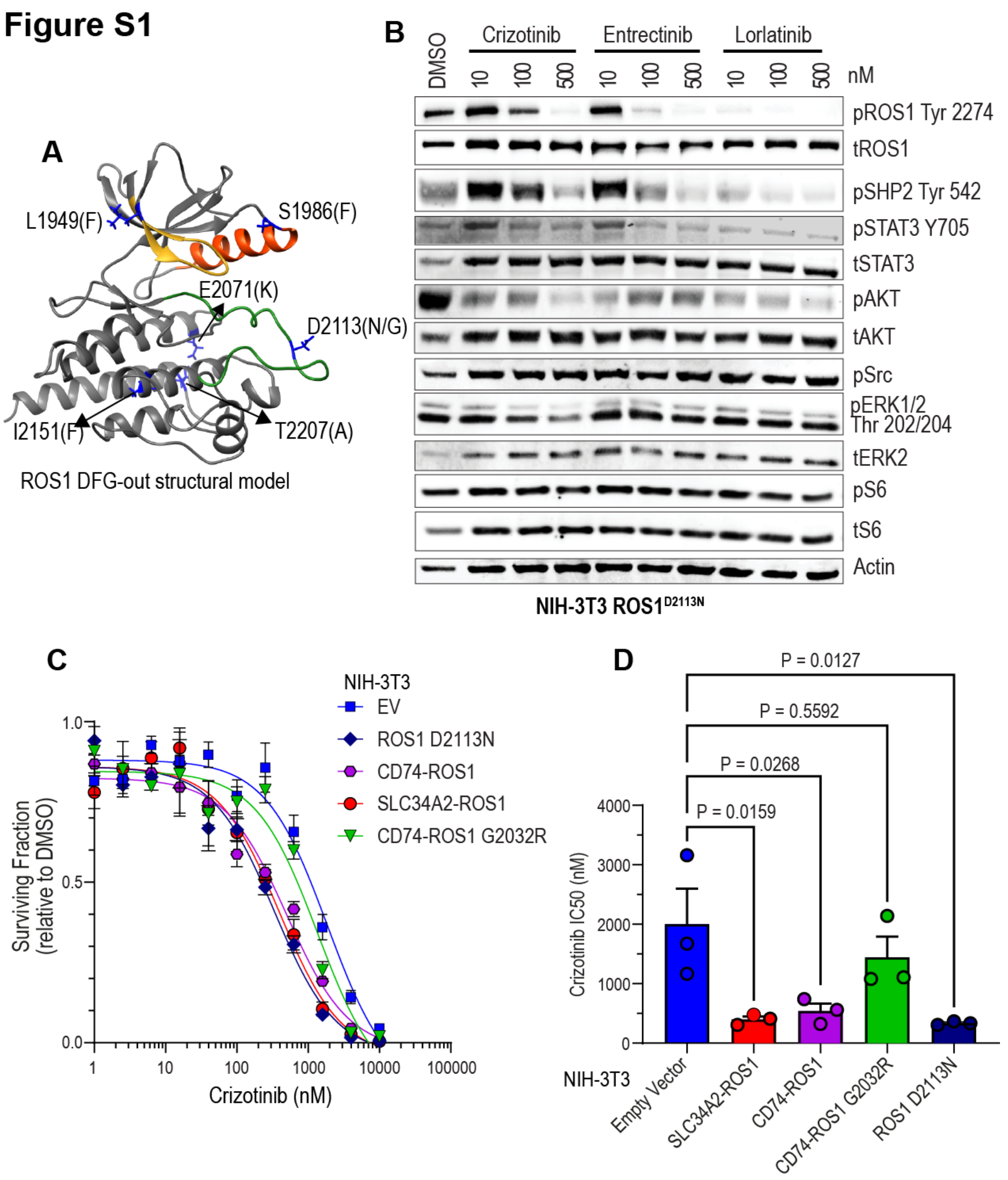


**Supplementary Figure S1. NIH-3T3 cells transduced with ROS1$^{D2113N}$ become sensitive to ROS1-TKI inhibition.** **A.** Ribbon diagram of ROS1 kinase domain structural model annotated with amino acid substitutions that increased catalytic activity. **B.** Immunoblot analysis of the phosphorylated (p) and total (t) ROS1 and effector proteins in NIH-3T3 ROS1$^{D2113N}$ cell lysates prepared after 4-hour treatment with vehicle (0.05% DMSO) or 10, 100, or 500 nM of crizotinib, entrectinib, or lorlatinib. **C.** Dose-response cell viability assay with NIH-3T3 ROS1$^{D2113N}$ cells treated with crizotinib (n = 3) as assessed by MTS colorimetric cell viability reagent. **D.** Bar graph of crizotinib $IC_{50}$ values from **(C)**. One-way ANOVA with Dunnett's multiple comparisons test used to assess statistical significance.

#### Figure S2

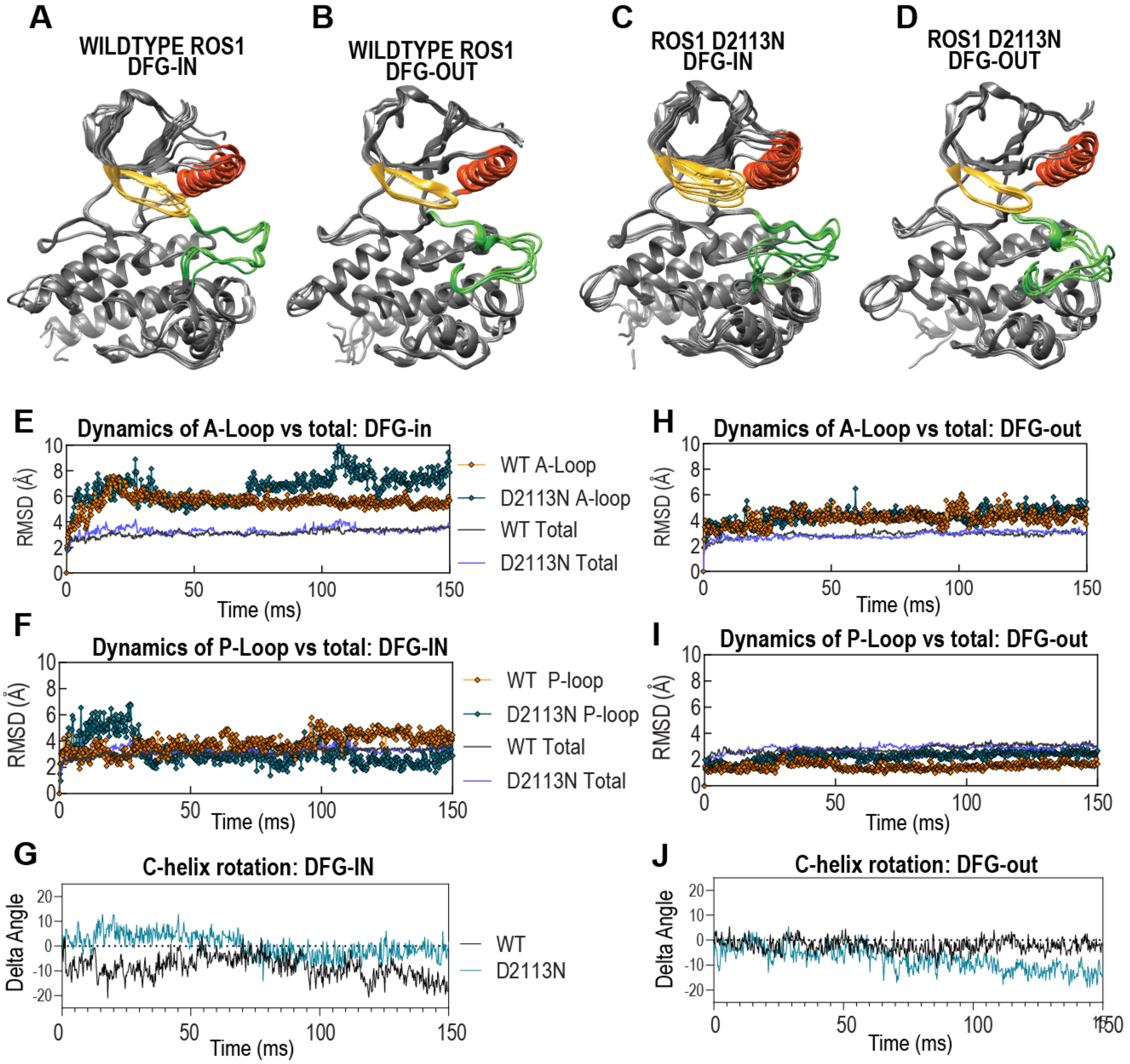


**Supplementary Figure S2. Molecular dynamic simulations of ROS1$^{WT}$ and ROS1$^{D2113N}$ kinase demonstrate effects on A-loop in the DFG-in conformation but minimal impact on other domains. A-D.** Six individual representative poses adopted by ROS1$^{WT}$ and ROS1$^{D2113N}$ in DFG-in and DFG-out conformations as indicated by labels. Root Mean Square Deviation values (Y-axis) plotted over 150 milliseconds of simulation duration (X-axis) for A-loop in DFG-in conformation (**E**), A-loop DFG-out conformation (**H**), P-loop in DFG-in (**F**) and DFG-out (**I**) conformations and αC-helix in DFG-in (**G**) and DFG-out (**J**) conformations.

**Figure S3**

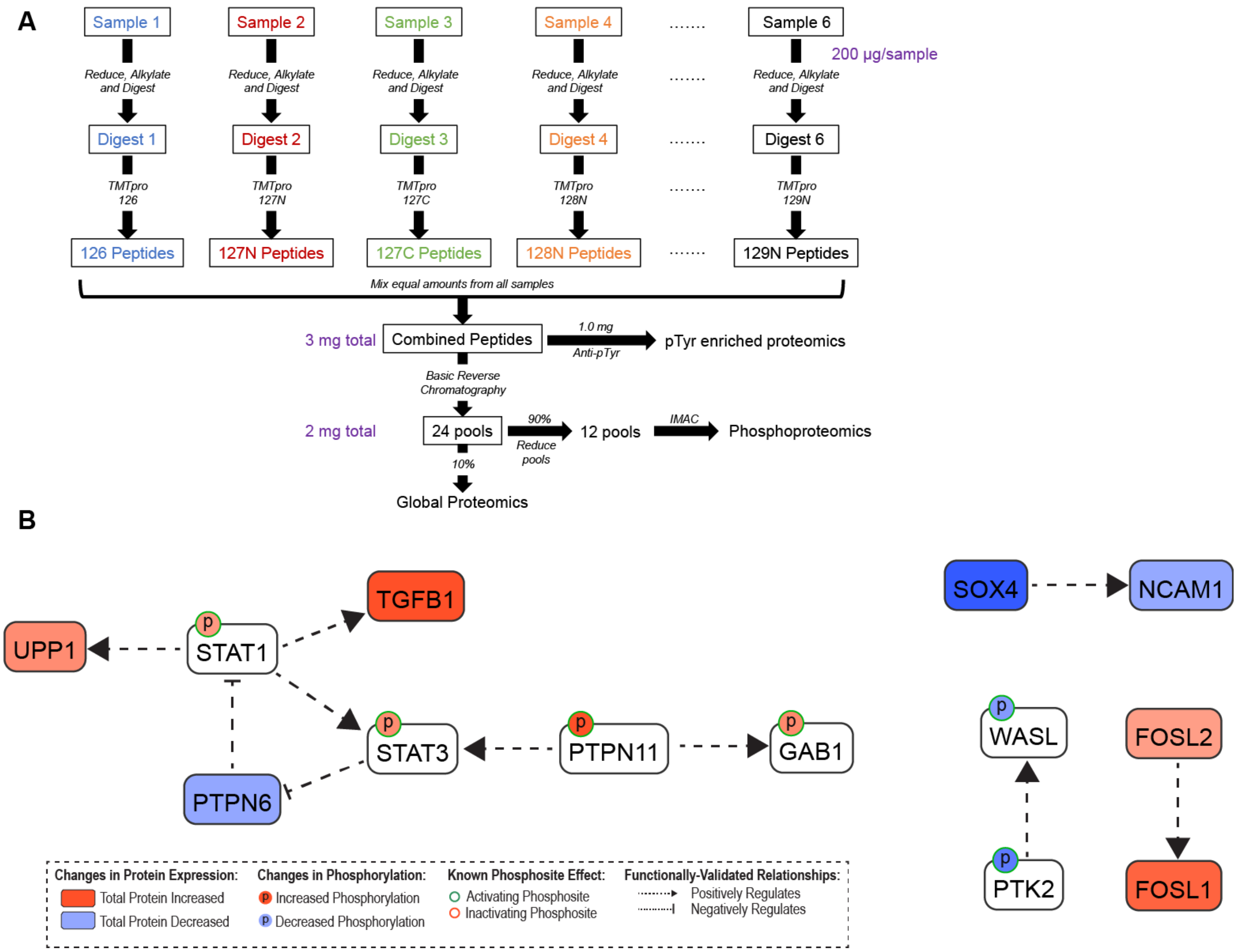


**Supplementary Figure S3: Workflow for global proteomics and phosphoproteomics enrichment and pathway analysis. A.** Setup of global proteomics and phosphoproteomics enrichment after lysate generation and quantification. **B.** Causalpath analysis of global proteomics and phosphoproteomics log2-transformed spectral counts highlights signaling changes promoted by ROS1$^{D2113N}$ relative to ROS1$^{WT}$.

### Figure S4

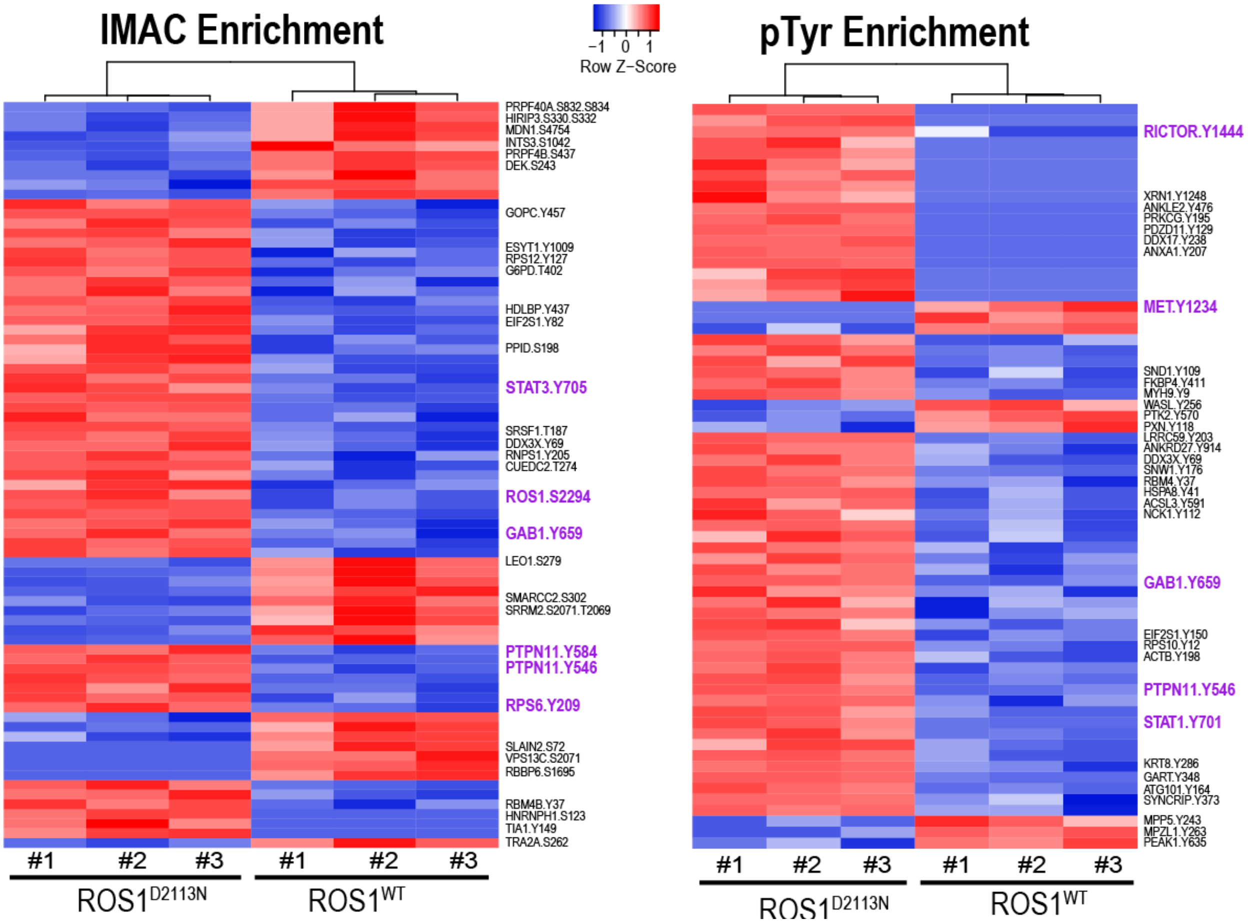


**Supplementary Figure S4: Differentially expressed proteins in ROS1$^{D2113N}$ relative to ROS1$^{WT}$ cells.** Heatmaps of IMAC enriched phosphoproteome (*left*) and phospho-tyrosine (pTyr) enriched phosphoproteome (*right*) of HEK-293A cells stably transduced with ROS1$^{WT}$ or ROS1$^{D2113N}$. Red color indicates upregulation and blue color indicates downregulation. All phosphosites displayed are statistically significant ($q \leq 0.1$ for IMAC; $q \leq 0.05$ for pTyr). Rows and columns were clustered with the Average Linking method while the distance measurement method was Euclidean.

#### Figure S5

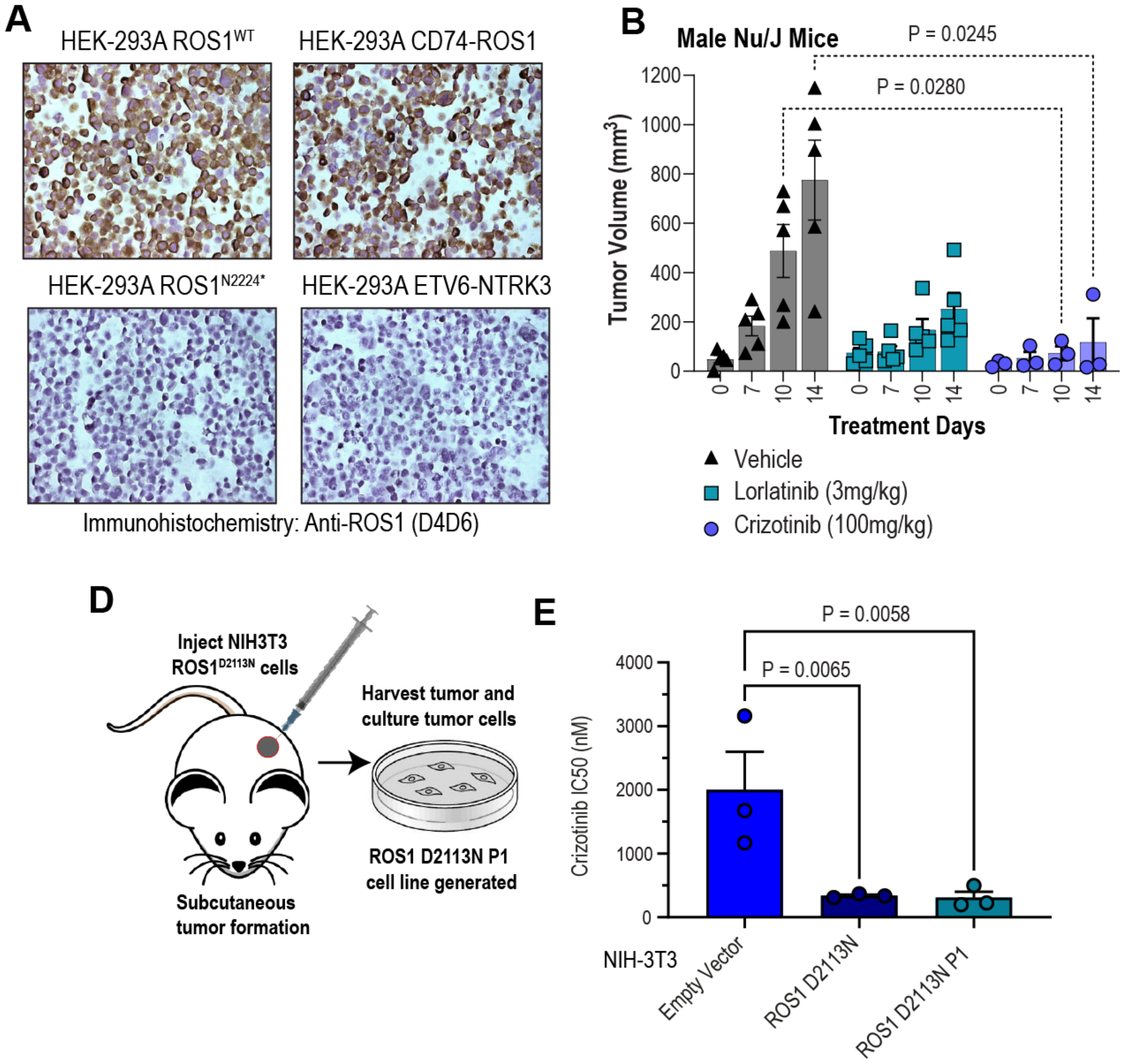


**Supplementary Figure S5. ROS1$^{D2113N}$ tumors and the tumor-derived cell line respond to ROS1-TKI treatment. A.** Immunohistochemistry (ICC) of 20µm sections from formalin-fixed, paraffin-embedded HEK-293A cell transduced with ROS1$^{WT}$, CD74-ROS1, ROS1$^{N2224*}$, and ETV6-NTRK3 to demonstrate specificity of ROS1 D4D6 antibody. **B.** Tumor growth of NIH-3T3 ROS1$^{D2113N}$ cells subcutaneously injected into male Nu/J mice and treated for 14 days with vehicle, crizotinib (100 mg/kg), or lorlatinib (3 mg/kg). n = 3. Two-way ANOVA with Dunnett's multiple comparisons test used to assess statistical significance. **C.** Diagram of generation of NIH-3T3 ROS1$^{D2113N}$ P1 cell line from subcutaneous tumor in Nu/J mice. **D.** Bar graph shows crizotinib $IC_{50}$ values from NIH-3T3 empty vector, ROS1$^{D2113N}$, and ROS1$^{D2113N}$ P1 cell lines as measured with MTS colorimetric cell viability reagent. n = 3. One-way ANOVA with Dunnett's multiple comparisons test used to assess statistical significance.

**Supplementary Movies – Legend**

**Movie S1.** MCF10A pCX4 (empty vector) live-cell imaging with confluence mask (blue outline)

**Movie S2.** MCF10A ROS1 wildtype (WT) live-cell imaging with confluence mask (blue outline)

**Movie S3.** MCF10A ROS1 D2113N live-cell imaging with confluence mask (blue outline)

**Movie S4.** MCF10A ROS1 D2113G live-cell imaging with confluence mask (blue outline)

**Movie S5.** MCF10A SLC34A2-ROS1 live-cell imaging with confluence mask (blue outline)
